# Pathobionts from chemically disrupted gut microbiota induce insulin-dependent diabetes in mice

**DOI:** 10.1101/2022.06.05.494898

**Authors:** Xin Yang, Zhiyi Wang, Junling Niu, Rui Zhai, Xinhe Xue, Guojun Wu, Guangxun Meng, Huijuan Yuan, Liping Zhao, Chenhong Zhang

## Abstract

**Background:** Dysbiotic gut microbiome, genetically predisposed or chemically disrupted, has been linked with insulin-dependent diabetes (IDD) including autoimmune type 1 diabetes (T1D) in both humans and animal models. However, specific IDD-inducing gut bacteria remain to be identified and their casual role in disease development demonstrated via experiments that can fulfill Koch’s postulates.

**Results:** Here, we show that novel gut pathobionts in the Muribaculaceae family, enriched by a low-dose dextran sulfate sodium (DSS) treatment, translocated to the pancreas and caused local inflammation, beta cell destruction and IDD in C57BL/6 mice. Antibiotic removal and transplantation of gut microbiota showed that this low DSS disrupted gut microbiota was both necessary and sufficient to induce IDD. Reduced butyrate content in the gut and decreased gene expression levels of an antimicrobial peptide in the pancreas allowed for the enrichment of members in the Muribaculaceae family in the gut and their translocation to the pancreas. Pure isolate of one such members induced IDD in wildtype germ-free mice on normal diet either alone or in combination with normal gut microbiome after gavaged into stomach and translocated to pancreas.

**Conclusion:** The pathobionts that are chemically enriched in dysbiotic gut microbiota are sufficient to induce insulin-dependent diabetes after translocation to the pancreas. This indicates that IDD can be mainly a microbiome-dependent disease, inspiring the need to search for novel pathobionts for IDD development in humans.

## Background

Insulin-dependent diabetes (IDD) refers to a wide range of diabetic conditions characterized by absolute insufficiency of insulin secretion [1]. The most prevalent type of IDD is type 1 diabetes (T1D), which is an autoimmune condition that leads to specific destruction of beta cells and complete loss of insulin-production [2]. Except autoimmune T1D in which genetics may play a critical role, IDD can be caused by environmental factors, such as treatment with immune checkpoint inhibitors, toxic compounds, and infection etc. [3–5]. The rapid increase in the incidence of IDD in the past three decades indicates that environmental factors, rather than genetics, may be the main driver of such diseases [6]. However, the pathological mechanism on how such environmental factors can induce IDD remain elusive.

It has been hypothesized that gut microbiota may work as a mediating factor in the development of IDD [6]. This hypothesis has been supported mainly by studies on the role of gut microbiota in the development of T1D in both animal models and human cohorts [7]. Altered gut microbiota in connection to T1D has been reported in Finnish, German, Italian, Mexican, American (Colorado) and Turkish children [8–10]. The longitudinal functional profiling of the developing gut microbiome of infants in TEDDY study showed that short-chain fatty acids (SCFAs) from fermentation of gut microbiota have protective effects in early-onset human T1D [8]. Studies have shown that the gut microbiota and its metabolites contribute to the development of T1D in NOD mice through regulating toll-like receptor-dependent signaling [11, 12], sex hormone-related pathways [13], IL-17 related mucosal immune reactions [14, 15], and pancreatic production of antimicrobial peptides [16]. However, germ-free mice derived from NOD mice can still develop T1D, indicating that the gut microbiota is not essential for the development of T1D in this animal model [13]. The development of IDD in mice treated with toxic compounds, such as streptozotocin (STZ), has also been associated with gut microbiota dysbiosis. Interestingly, the removal of the gut microbiota by an antibiotic cocktail blocked IDD induction by STZ, showing that the gut microbiota may be indispensable for this chemically-induced IDD [7]. However, specific members of the dysbiotic gut microbiota which may drive IDD, and the associated etiological mechanisms remain to be elucidated.

Many toxic compounds have been shown to disrupt the gut microbiota and aggravate inflammation-related diseases, including metabolic diseases [17]. Notably, dextran sulfate sodium (DSS), a compound with very low bioavailability, has been well known for its effect on disrupting the gut microbiota and inducing acute colitis at a high dose (2-3%) and chronic colitis at a low dose (0.5-1%) when administered via drinking water in mice [18–20]. To explore how an even lower level of DSS would affect the gut microbiota and host health, we reduced the dose to 0.2%. We serendipitously found that the mice did not show inflammation or impaired barrier function in their gut; instead, they developed IDD with pancreatic beta cells significantly destructed. We identified and isolated one of the pathobionts in family Muribaculaceae which was enriched in the low DSS-disrupted gut microbiota and confirmed that this single pathobiont is sufficient to induce the initiation and progression of IDD in germfree mice. This study indicates that IDD can be a microbiome-dependent disease, inspiring the search of such novel pathobionts for IDD development in humans.

## Methods

### Animal Trials

#### Mice

Five-week-old SPF male mice (C57BL/6) were obtained from SLAC Inc. (Shanghai, China) and kept under SPF environment at the animal facility of Shanghai Jiao Tong University, Shanghai, China. Mice (five mice/cage) were housed in IVC system cages (Dimensions 325 × 210 × 180 mm; Suzhou Fengshi Laboratory Animal Equipment Co., Ltd). The mice were fed with a sterilized normal chow diet (10% energy from fat; 3.25 kcal/g; 1010009, SLAC) and were housed in a room maintained at 22.0 ± 1°C with 30-70% humidity, with a 12 h light/dark phase cycle (lights on at 07:00 am).

Eight-week-old of germ-free male C57BL/6 mice were maintained in flexible-film plastic isolators at the Laboratory Animal Center of the SLAC Inc. (Shanghai, China). The mice were fed a sterilized normal chow diet (10% energy from fat; 3.25 kcal/g; P1103F-50, SLAC). The mice were housed in a room maintained at 22.0 ± 1°C with 30-70% humidity with a 12 h light/dark phase cycle (lights on at 06:00 am). Surveillance for bacterial contamination was performed by periodic bacteriological examinations of feces, food and padding. Detailed operations of surveillance for bacterial contamination were consistent with previous work [21].

#### Animal Trials

**Trial-1:** After one-week of acclimation, the mice were randomly assigned to 2 groups: (i) NC group (n = 10), control mice provided pure drinking water for feeding 12 weeks; (ii) DSS group (n = 10), mice treated with 0.2% (w/v) DSS (0216011080, MP Biomedicals, USA) to drinking water for feeding 12 weeks. Independently repeated trials were performed three times.

**Trial-2:** After one-week of acclimation, a cocktail of antibiotics (0.5 g/L of vancomycin, 1 g/L of ampicillin, 1 g/L of neomycin, 1 g/L of metronidazole) was introduced in drinking water [22]. For 5 weeks. The mice were then randomly assigned to 2 groups: (i) ABX group (n = 10), antibiotic cocktail-treated mice provided pure drinking water for 12 weeks; (ii) ABX+DSS group (n = 8), antibiotic cocktail-treated mice with 0.2% (w/v) DSS added to drinking water for 12 weeks. During the 12 weeks, both groups continued to have the antibiotic cocktail in their drinking water.

**Trial-3:** After one-week of acclimation, the antibiotic cocktail was introduced in drinking water for 5 weeks. The mice were then randomly assigned to 2 groups: (i) FM*_NC_* group (n = 10), antibiotic cocktail-treated mice inoculated with the fecal inoculum from mice in the NC group and provided pure water for feeding 20 days; (ii) FM*_DSS_* group (n = 10), antibiotic cocktail-treated mice inoculated with the fecal inoculum from mice in the DSS group and provided pure water for feeding 20 days. The oral gavage was performed once a day for 3 times (Day 0, Day 1 and Day 4). Each mouse was given 200 µL of inoculum every time.

**Trial-4:** After one-week of acclimation, the antibiotic cocktail was introduced in drinking water for 5 weeks. The mice were then randomly assigned to 2 groups: (i) FM*_NC_* group (n = 10), antibiotic cocktail-treated mice inoculated with the fecal inoculum from mice in the NC group and provided pure drinking water for 20 days; (ii) FM*_NC+Muri_* group (n = 10), antibiotic cocktail-treated mice inoculated with a mixture of the fecal inoculum of the NC group and the MF 13079 strain and provided pure drinking water for 20 days. The oral gavage was performed once a day for 3 times (Day 0, Day 1 and Day 4). Each mouse was given 200 µL of fecal inoculum every time in FM*_NC_* group; and each mouse was given 100 µL fecal inoculum and 100 µL bacterial suspension in FM*_NC+Muri_* group.

**Trial-5:** The germ-free mice were randomly assigned to two groups and each group was kept in an individual plastic isolator. (i) Akk group (n = 12), germ-free mice inoculated with the *Akkermansia muciniphila* strain 139 and provided pure drinking water for 14 days; (ii) Muri group (n = 11), germ-free mice inoculated with the MF 13079 strain and provided pure drinking water for 14 days. The oral gavage was performed once a day for 2 consecutive days. Each mouse was given 200 µL of bacterial suspension every time.

### Preparation of Inoculum

#### Preparation of inoculum from mice feces

Fresh fecal samples from each mouse in NC and DSS groups (in the 12th week of Trial-1) were collected. The fecal samples in the same group were mixed in anaerobic sterile Ringer working buffer (9 g/L of sodium chloride, 0.4 g/L of potassium chloride, 0.25 g/L of calcium chloride dehydrate and 0.05% (w/v) L-cysteine hydrochloride) in an anaerobic chamber (80% N_2_ :10% CO_2_ :10% H_2_) [23], and diluted to 50 times. Then the suspensions were added with 20% (w/v) skim milk (LP0031, Oxoid, UK) and the final fecal suspensions were diluted to 100 times. The fecal suspensions were preserved at − 80 °C until inoculation.

#### Preparation of bacterial suspensions

*Akkermansia muciniphila* strain 139 was cultured in a synthetic medium [24] and Muribaculaceae strain MF13079 was cultured in a modified MPYG medium (Table S1). The harvested bacterial cells were washed twice with PBS (pH 7.4) and resuspended in PBS (pH 7.4) and added with equal volume of 20% skim milk to a density of 10^9^ cells/ml. The bacterial suspensions were stored at − 80 °C until inoculation.

### Glucose Tolerance Test and Estimation of Insulin Sensitivity

#### Oral glucose tolerance test (OGTT)

Mice were fasted for 6 h and baseline blood glucose levels were measured with an Accu-Check Performa blood glucose meter (Roche, USA) using blood collected from the tail vein. After administration of glucose (2.0 g/kg body weight) via oral gavage, blood glucose levels were measured at 15, 30, 60, and 120 min. Serum insulin levels were also measured by ELISA kit (90080, Crystal Chem, USA) at 0, 15, 30, 60 min after glucose challenge.

#### Insulin tolerance test (ITT)

Six hours fasted mice were injected intraperitoneally with 1.0 U insulin/kg body weight (Novolin, Novo Alle, Denmark). Blood glucose was then measured at 0, 15, 30, 60, and 120 min after injection with the blood glucose meter (Roche, USA).

### *In vivo* Intestinal Permeability Assay

Intestinal permeability was measured based on the permeability to 4000 Da-FITC-dextran in plasma (DX-4000-FITC) (46944; Sigma-Aldrich, St. Louis, Missouri, USA) as previously described [25]. Briefly, after fasting for 6 h, the mice were administered with DX-4000-FITC (500 mg/kg body weight, 125 mg/mL) by oral gavage. After 4 h, blood plasma collected from the tip of the tail vein was diluted with equal volume of PBS (pH 7.4) and analyzed for DX-4000-FITC concentration with a fluorescence spectrophotometer (HTS-7000 Plus-plate-reader; Perkin Elmer, Wellesley, Massachusetts, USA) at an excitation wavelength of 485 nm and emission wavelength of 535 nm. A standard curve was obtained by diluting FITC-dextran in non-treated plasma diluted with PBS (1:3 v/v).

### Energy Assessment of Feces

#### Fecal output measurement

Mice were placed in a clean cage individually and feces were collected for 24 h. Dry weights were taken after fecal pellets were incubated at 65°C in an oven for 24 h (JC101, Shanghai Chengshun Instrument & Meter Co., Ltd, Shanghai, China).

#### Bomb calorimetry

Bomb calorimetry was performed on dried fecal pellets. Gross energy content was measured using an isoperibol bomb calorimeter (C6000, IKA, Germany). The calorimeter energy equivalent factor was determined using benzoic acid standards.

### Triglycerides and Cholesterol Measurement

Serum cholesterol and triglycerides were quantified with colorimetric kits (A111-1-1-1 and A110-1-1, respectively) from Nanjing Jiancheng Bioengineering Institute (Nanjing, China).

Frozen liver sample was homogenized in a corresponding volume (1:9, w:v) of homogenizing buffer (pH 7.4, 0.01 mol/L Tris-HCl, 0.1 mmol/L EDTA-2Na, 0.8% NaCl). The supernatant was collected after centrifugation for 25 min at 2,000×*g* and 4 °C. Cholesterol and triglycerides in homogenized tissue were quantified by colorimetric kits from Nanjing Jiancheng Bioengineering Institute (Nanjing, China). Tissue homogenized total protein was measured with the Protein Quantification Kit (20202ES76; Yeasen Biotechnology Ltd., Shanghai, China).

### Amylase and Lipase Measurement

Serum amylase and lipase were quantified with colorimetric kits (C016-1-1 and A054-2-1, respectively) from Nanjing Jiancheng Bioengineering Institute (Nanjing, China).

### Fecal Lipocalin-2 Detection

Fecal lipocalin-2 content was quantified with a Lipocalin-2/NGAL Quantikine ELISA kit (DLCN20, R&D Systems).

### mRNA Expression of Genes Related with Gut Barrier Integrity in the Colon

The total RNA of colons was extracted using the RNeasy Plus Universal Tissue Mini Kit (73404, Qiagen, Duesseldorf, Germany). The RNA was reverse-transcribed to cDNA using the SuperScript™ III First-Strand Synthesis (18080051, Invitrogen, Calif., USA). RT-qPCR was performed with the Light Cycler 96 (Roche, Geneva, Switzerland) using iQ SYBR Green Supermix (170-8882AP, BIO-RAD, CA, USA). The PCR conditions were 95°C for 3 min, followed by 40 cycles of 95°C for 20 s, 56°C for 30 s, and 72°C for 30 s, and plate reads for 5 s. Gene expression levels were determined using the comparative ΔΔC*_T_* method (2^−ΔΔC^*_T_* method), with the β-actin gene serving as the housekeeping gene. Forward (F) and reverse (R) primer sequences are as follows:

*β-actin*, F-CCTTCTTGGGTATGGAATCCTGTG, and R-CAGCACTGTGTTGGCATAGAGG;

*Tjp*1, F-ACCCGAAACTGATGCTGTGGATAG, and R-AAATGGCCGGGCAGAACTTGTGTA;

*Cldn*1, F-GATGTGGATGGCTGTCATTG, and R-CCTGGCCAAATTCATACCTG; *Muc*1, F-TACCCTACCTACCACACTCACG, and R-CTGCTACTGCCATTACCTGC;

*Muc*2, F-CACCAACACGTCAAAAATCG, and R-GGTCTCTCGATCACCACCAT; *Muc*3, F-CTTCCAGCCTTCCCTAAACC, and R-TCCACAGATCCATGCAAAAC; *Muc*4, F-GAGAGTTCCCTGGCTGTGTC, and R-GGACATGGGTGTCTGTGTTG;

*Zo*-1, F-ACCCGAAACTGATGCTGTGGATAG, and R-AAATGGCCGGGCAGAACTTGTGTA;

*Occludin*, F-ATGTCCGGCCGATGCTCTC, and R-TTTGGCTGCTCTTGGGTCTGTAT.

### Histopathology

Fresh epididymal fat pads were fixed with 4% paraformaldehyde for 48 h, embedded in paraffin, sectioned, and stained with hematoxylin and eosin (H&E) (G1003; Wuhan Servicebio technology Ltd., Wuhan, China). Digital images of sections were acquired with a Leica DMRBE microscope. Adipocyte sizes were assessed using Image Pro Plus v6.0 (Media Cybernetics Inc., Silver Springs, MD, USA). For each mouse, mean areas of adipocytes were determined in five discontinuous scans under ×200 magnification.

Fresh liver tissues were fixed with 4% paraformaldehyde for 48 h, embedded in paraffin, sectioned, and stained with hematoxylin and eosin (H&E) (G1003; Wuhan Servicebio technology Ltd., Wuhan, China). Digital images of sections were acquired with a Leica DMRBE microscope. For Oil Red O staining of liver sections, liver tissues were fixed in 4% paraformaldehyde for at least 48 h, embedded in optimal cutting temperature (OCT) compound (4583; CellPath Ltd., Newtown, Powys, UK) and frozen at −20 °C. Sections were stained with Oil Red O (O0625; Sigma-Aldrich Ltd., St. Louis, MO, USA). Digital images of sections were acquired with a Leica DMRBE microscope.

Fresh pancreatic tissues were fixed with 4% paraformaldehyde for 48 h, embedded in paraffin, sectioned, and stained with hematoxylin and eosin (H&E) (G1003; Wuhan Servicebio technology Ltd., Wuhan, China). Digital images of sections were acquired with a Leica DMRBE microscope. pancreas islet number were assessed using Image Pro Plus v6.0 (Media Cybernetics Inc., Silver Springs, MD, USA). Histological analysis of pancreatitis was determined by one observer who was blinded to the treatment group. Scores of pancreatitis severity was assessed as described previously[26].

### Immunofluorescence Observation of Insulin in the Pancreas

Fresh pancreatic tissues were fixed with 4% paraformaldehyde and then embedded in paraffin. Subsequently, 4 μm sections were blocked with 5% bovine albumin (BSA), and stained overnight with anti-insulin pAb (1:500, Abcam, ab7842) at 4°C. After washing, anti-pig-AlexaFluor488 secondary antibody as the second-step reagents were applied. The sections were incubated for 15 min at room temperature in the dark. Nuclei were stained with DAPI (G1012, Servicebio). Digital images of sections were acquired with a Leica DMRBE microscope after completely rinsing with sterile PBS. The insulin-positive area of each section was counted with Image Pro Plus v6.0 (Media Cybernetics Inc., Silver Springs, MD, USA).

### Immunohistochemical Analysis of ZO-1 and Occludin in the Colon

After deparaffinization and processing for antigen retrieval, the blank colon sections were incubated overnight at room temperature with anti-rabbit ZO-1 antibody (1:500, catalog no. GB11195; Servicebio, China) and Occludin antibody (1:200, catalog no. GB11149; Servicebio) separately. Sections were then incubated with anti-rabbit IgG secondary antibody (KPL, MA, USA) for 50 min at room temperature. After washing with PBS, the DAB Envision kit (catalog no. K5007; Dako, Copenhagen, Denmark) was used to develop color; ZO-1 and Occludin appear brown, while nuclei are blue. Images of each colon section were obtained in tripartite by experienced staff who were blind to the experiment under ×200 magnification using a Leica DMRBE microscope and were analyzed using Image Pro Plus 6.0, according to the method previously described [27]. The integrated optical density (IOD) values were log10 transformed.

### Flow Cytometry Evaluation of Subtypes of Leukocytes in the Pancreas

Fresh pre-cut pancreatic tissue (2 × 2 mm) were digested with a solution of collagenase P in HBSS-1% HEPES (0.75 mg/mL, Roche) at 37°C for 7 min with shaking. Digestion was stopped by adding HBSS-10% FCS 1% EDTA followed by extensive washes by FACS buffer (1×PBS pH7.2, 0.5% BSA, 2mM EDTA). The suspensions were filtered through 100 μm cell strainer (352360, FALCON, USA). Suspensions were transfer into Polystyrene Round-Bottom Tube (352052, FALCON, USA) followed by centrifugation at 400×g at 4℃ for 5 min. To ensure single cell bioactivity, it was necessary to disperse the cells quickly after discarding the supernatant. 500 μL FACS buffer was added to re-suspend cells. The wash step was repeated twice to get a single cell suspension. Non-specific binding was blocked with FcR blocking antibody (anti-mouse CD16/32, 553142, BD Pharmingen) before surface staining with the following mAbs: Anti-mouse CD45.2 (563051, BD Horizon), Anti-mouse-CD3e (562600, BD Horizon), Anti-mouse-CD4 (553653, BD Pharmingen), Anti-mouse-CD8a (551162, BD Pharmingen), Anti-mouse-F4/80 (12-4801-82, eBioscience). Single cell suspension was washed twice as described above by 500 μL FACS buffer. Intracellular staining was carried out after fixation and permeabilization (00-5523-00, eBioscience) with the following antibodies: Anti-mouse Foxp3 (77-5775-40, eBioscience), Anti-mouse-IL-17A (12-7177-81, eBioscience). Dead cells were excluded with a fixable viability dye (65-086514, Invitrogen, USA). The fluorescence was examined on an Flowcytometer (BD LSRFortessa, Becton, Dickinson and Company, USA).

### Sequence-guided Bacteria Isolation

Fresh fecal samples from DSS group were collected and mixed in anaerobic sterile Ringer working buffer in an anaerobic workstation (Don Whitley Scientific Ltd, Shipley, UK). The 10^5^ diluted suspension with Ringer working buffer was placed onto 46 medium plates (Table S1) and incubated under anaerobic condition (80% N_2_, 10% CO_2_, and 10% H_2_) at 37°C for 7 days. The 16S rRNA gene of each colony was obtained using the primers 27-F (AGAGTTTGATCCTGGCTCAG) and 1492-R (GGTTACCTTGTTACGACTT). The obtained 16S rRNA sequences were sequenced and aligned with the ASVs in Muribaculaceae enriched in the gut of DSS-treated mice. One Muribaculaceae strain (MF13079) was isolated using a previously reported MPYG medium [28] which was modified by adding 3% fetal bovine serum (FBS, GIBCO, 10099-141). The Muribaculaceae strain MF13079 16S rRNA sequence was aligned with the GenBank. All the 16S rRNA sequence of the isolated strains in the family Muribaculaceae and the 16S rRNA sequence of the representative strains in other families of the phylum Bacteroidetes were selected to build a phylogenetic tree by using the Neighbor-Joining method in MEGA 6.

### Whole-Genome Sequencing and Analysis of MF13079

Whole-genome DNA of Muribaculaceae strain MF13079 was extracted with QIAamp BiOstic Bacteremia DNA Kit (12240-50, Qiagen, USA) and sequenced with the Oxford^@^ Nanopore PromethION platform (performed by Nextomics Biosciences, Wuhan, China). Subreads of the sequence were assembled into a single complete chromosome using the HGAP 2.3.0 pipeline [29].

The genome sequence of Muribaculaceae strain MF13079 was adjusted to start with the replicational origin (location of *DnaA*) for comparison. Protein-coding sequences (CDSs), tRNAs, and rRNAs were predicted and annotated using the Prokka 1.12 pipeline (Table S2) [30]. Functional annotation of genes related with flagella-synthesis in the CDSs of MF13079 was performed using the Kyoto Encyclopedia of Genes and Genomes (KEGG) database.

### Microbiota Profiling in Tissues

#### DNA extraction of microorganisms in tissues

Pre-cut pancreatic tissue (5 × 5 mm) and mesenteric lymphatic nodes (MLNs) were homogenized using a sterile pestle in a sterile mortar under liquid nitrogen as the media. Homogenate was used for DNA extraction by QIAamp Pro DNA Kit (51804, Qiagen, Duesseldorf, Germany). Sterile DNA-free water was used as negative controls.

#### Bacterial load in the pancreas and mesenteric lymphatic nodes (MLNs)

Nested PCR was used in this situation. The 16S rRNA gene was first amplified using the primers 27-F (AGAGTTTGATCCTGGCTCAG) and 1492-R (GGTTACCTTGTTACGACTT). PCR was performed in a volume of 25.0 μl with 10 ng DNA template, 1 μl of each primer (12.5 μM), 2.5 μl of 10× PCR buffer, 0.3 μl of Taq polymerase (TaKaRa), 1 μl of dNTP mix (2.5 mM). The reaction conditions consisted of initial denaturation at 94 °C for 5 min; 25 cycles of 94 °C for 30 s, 55 °C for 30 s and 72 °C for 90 s; and a final extension step for 10 min at 72 °C. Subsequently, a 466 bp length sequence located in 16S rRNA gene was used for quantitative PCR through the primers Uni331F (TCCTACGGGAGGCAGCAGT) and Uni797R (GGACTACCAGGGTATCTAATCCTG TT) [31]. Briefly, qPCR was performed with the Light Cycler 96 (Roche, Geneva, Switzerland) using iQ SYBR Green Supermix (170-8882AP, BIO-RAD, CA, USA). The PCR conditions were 95°C for 3 min, followed by 40 cycles of 95°C for 15 s, 60°C for 60 s and plate read for 5 s at 80°C. A whole-length 16S rRNA gene of an *Akkermansia muciniphila* strain derived from mice intestine was used to plot a standard curve to calculate the copies of 16S rRNA genes in the samples.

#### Fluorescence in situ hybridization (FISH) of 16S rRNA in the pancreas

The EUB338 16S rRNA probe labeled with the fluorophore Cy3 (extinction wavelength, 555 nm; emission wavelength, 570 nm; Servicebio, China) was used to detect the bacterial colonization within mouse pancreatic tissues by FISH. Nuclei were stained with DAPI (G1012, Servicebio). Fluorescence microscopic analysis was conducted with Nikon Eclipse 90i confocal microscope (Nikon, Melville, NY) using a Cy3 labeled-probe at 1μM as described [32–34].

#### Pancreatic microbiota analysis by sequencing of V3-V4 region in 16S rRNA gene

The 16S rRNA gene was first amplified from total DNA of pancreatic tissue using the primers 27-F (AGAGTTTGATCCTGGCTCAG) and 1492-R (GGTTACCTTGTTACGACTT). The PCR products were used as a template, a sequencing library of the V3-V4 region of the 16S rRNA gene was constructed following the manufacturer’s instructions (Part # 15044223Rev.B; Illumina Inc., San Diego, CA, USA) with improvement as previously described [20], and sequenced on the Illumina MiSeq platform (Illumina, Inc., San Diego, CA, USA). The negative controls from DNA extraction step were amplified and sequenced. Sterile DNA-free water was also used as template for PCR in sequencing library preparation and then sequenced.

The raw paired-end reads were processed and analyzed using the Quantitative Insights into Microbial Ecology 2 (QIIME2, v2019.01) platform [28]. Demultiplexed sequence data was imported into QIIME2, adapters and primers were trimmed. Amplicon sequence variants (ASVs) from each sample was inferred by using the DADA2 pipeline [35] for filtering, dereplication, sample inference, merging of paired-end reads and chimera identification. In the process of running the DADA2 pipeline, based on the quality profile of the data, forward and reverse reads were trimmed accordingly to ensure that the median quality score for each position is above 32. Potential reagent and environmental contaminations were identified using “decontam (v.1.2.1)” package in R (v3.4.4) based on the frequency of the ASV in the negative controls with threshold set at 0.5 [36]. The identified contaminants were removed from dataset. After removing contaminants, all samples were rarefied to 6,000 reads per sample for downstream analysis.

### Gut Microbiota Profiling

The bacterial DNA extraction from fecal samples of mice was performed as previously described [37]. A sequencing library of the V3-V4 region of the 16S rRNA gene was constructed following the manufacturer’s instructions (Part # 15044223Rev.B; Illumina Inc., San Diego, CA, USA) with improvement as previously described [20], and sequenced on the Illumina MiSeq platform (Illumina, Inc., San Diego, CA, USA).

The raw paired-end reads were processed and analyzed using the QIIME2. Demultiplexed sequence data was imported into QIIME2, adapters and primers were trimmed. ASVs from each sample was inferred by using the DADA2 pipeline for filtering, dereplication, sample inference, merging of paired-end reads and chimera identification. In the process of running the DADA2 pipeline, based on the quality profile of the data, forward and reverse reads were trimmed accordingly to ensure that the median quality score for each position is above 32. The taxonomy of all ASVs were annotated by SILVA (v132) reference database [38]. All samples were rarefied to 10,000 reads per sample for downstream analysis. Principal coordinate analysis (PCoA) of ASVs based on Bray-Curtis distance was performed using QIIME2. Subsequently the statistical significance was assessed using permutational multivariate analysis of variance (PERMANOVA) with 9,999 permutations and *P* values were adjusted for multiple comparison with Benjamini-Hochberg method [39].

Sparse partial least-squares discriminant analysis (sPLS-DA) models [40] were established to identify specific ASVs that contributed to the segregation of gut microbial structure in NC and DSS mice using “mixOmics (v6.3.1)” package [41] in R (v3.4.4). Centered log ratio (CLR) transformations of the relative abundance of ASVs were implemented in sPLS-DA models. The optimal classification performances of the sPLS-DA models were estimated by the perf function using 5-fold cross-validation with the smallest balanced error rate.

### Statistical analysis

Data in point plots and bar plots, horizontal lines indicate Mean ± S.E.M. Statistical significance was assessed by Student’s two-tailed t test using GraphPad Prism (version 8), with *P* values of <0.05 considered significant.

## Results

### Low-dose DSS induced gut microbiota-dependent IDD

To determine how a lower than usual dose of DSS would affect gut microbiota and host health, we randomized 6-week-old C57BL/6 mice into a treatment group with 0.2% DSS in the drinking water or a control group with no any DSS intake. Compared with control mice, the mice that received 12 weeks of low dose DSS treatment showed a significant increase in blood glucose levels and decrease in insulin levels in oral glucose tolerance test (OGTT) (Figure 1A and 1B). Moreover, the low dose DSS-treated mice and control did not differ in blood glucose levels in an insulin tolerance test (ITT), indicating that the low dose DSS-treated group did not develop insulin resistance (Figure 1C). Compared with the control mice, low dose DSS-treated mice had significantly decreased insulin expression in the pancreas (Figure 1D). We also found that the fasting C-peptide level in serum was significantly lower in mice with low dose DSS treatment than the control mice, demonstrating that the abnormal high glucose level was due to low insulin production and secretion (Figure S1). The low dose DSS-treated mice showed islet destruction in the pancreas (Figure 1E). These mice also showed mild pancreatitis, as indicated by histological characteristics, such as inflammatory cell infiltration and edema in the pancreas (Figure 1F), and an increase in serum amylase levels (Figure S2). The DSS-treated mice had significantly higher daily food and water intake than the control mice starting from the 8^th^ week of treatment (Figure S3). However, the energy consumption, body weight gain, and lipid metabolism did not differ between low dose DSS treatment and control throughout the study period (Figures S4). Thus, the treatment with 0.2% DSS in drinking water induced IDD in wild-type C57BL/6 mice.

**Figure 1.**
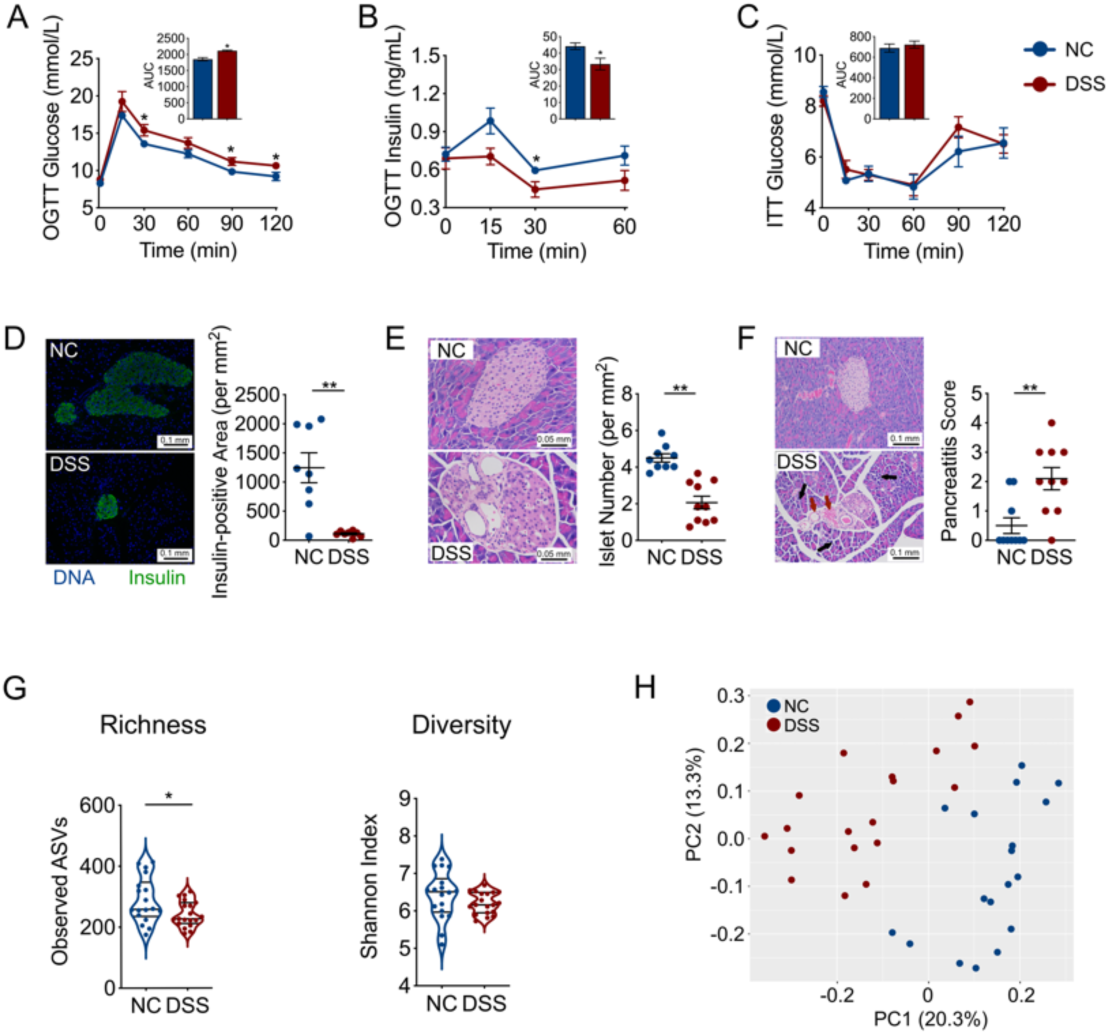
IDD induced by 0.2% DSS treatment in mice. **(A)** Blood glucose and **(B)** insulin levels of normal control (NC) and DSS-treated mice, as measured by an oral glucose tolerance test (OGTT) (the area under the curve (AUC) is shown in the inset panels). NC group, n=9; DSS group, n=7. **(C)** Blood glucose levels of NC and DSS-treated mice as measured by an insulin tolerance test (ITT) (the AUC is shown in the inset panels). NC group, n=8; DSS group, n=9. **(D)** Histological sections of pancreatic tissue (200×, scale bar = 0.1 mm) from NC and DSS-treated mice stained for insulin (green) and DNA (blue), and the insulin-positive area was quantified (at right). NC group, n=8; DSS group, n=8. **(E)** The number of islets per mm^2^ of pancreatic tissue was calculated based on H&E-stained histological sections of pancreatic tissue (400×, scale bar = 0.05 mm) from NC and DSS-treated mice. NC group, n=9; DSS group, n=10. **(F)** The histologic pancreatitis score was evaluated by edema, inflammation, and vacuolization in the H&E-stained histological sections of pancreatic tissue (200×, scale bar = 0.1 mm) from NC and DSS-treated mice. Black arrows, scattered structure; Red arrows, increased peri-islet neovascularization. NC group, n=10; DSS group, n=10. **(G)** The alpha-diversity of gut microbiota in NC and DSS-treated mice. **(H)** Overall gut microbial structure in NC and DSS-treated mice. Principal coordinate analysis (PCoA) was performed on the basis of Bray-Curtis distance at the amplicon sequence variant (ASV) level. For **(G)** and **(H)**, NC group, n=17; DSS group, n=18. The data in **(A)-(F)** are shown as the mean ± SEM, and Student’s t-test (two-tailed) was used to analyze difference between NC and DSS groups. \**P* <0.05 and \*\**P* <0.01. NC, control mice provided pure drinking water; DSS, mice treated with 0.2% (w/v) DSS in drinking water.

However, unlike higher doses DSS [42], this low dose DSS treatment did not induce colitis or gut barrier impairment in mice. H&E-stained histological sections of colon tissue showed that there were no structural changes of epithelium in the mice with 0.2% DSS in drinking water compared with the control (Figure S5). The fluorescein isothiocyanate (FITC)-dextran test, levels of mRNA expression of 8 different genes related with gut barrier integrity and immunohistochemical staining of Occludin and Zo-1 showed no significant difference in intestinal permeability in mice with or without 0.2% DSS treatment (Figure S5). Moreover, the fecal lipocalin-2 content, a biomarker for gut inflammation, had no significant difference between the 0.2% DSS-treated and control mice (Figure S5).

Although the dose of DSS used in our model was much lower than the dose needed to induce colitis [18], the gut microbiota structure in the low dose DSS-treated mice was significantly shifted away from the control, as shown by its reduced richness (Figure 1G) and significant difference in principal coordinate analysis (PCoA) score plot of Bray-Curtis distance based on the amplicon sequence variants (ASVs) of the 16S rRNA gene V3-V4 region (*P*=0.001 by permutational multivariate analysis of variance (PERMANOVA) test with 999 permutations, Figure 1H). Thus, the development of low dose DSS-induced IDD in mice was accompanied by significant changes in the gut microbiota without inducing colitis.

To find out whether the disrupted gut microbiota was necessary for the development of IDD induced by low dose DSS, we treated the adult C57BL/6 mice with a cocktail of antibiotics for 5 weeks that depleted the gut microbiota by more than 99% (Figure S6). We then introduced 0.2% DSS into the drinking water of the mice which were continuously treated with the antibiotic cocktail. Under the gut microbiota-depleted condition, DSS-treated (ABX+DSS) and control (ABX) mice had indistinguishable glucose metabolism, insulin secretion and histology in the pancreas (Figure 2, Figure S2). Thus, the removal of the gut microbiota abolished the low dose DSS-induced IDD in mice, indicating that the gut microbiota is essential for the disease induction. To test if the DSS-disrupted gut microbiota is sufficient to induce IDD, we transferred feces from control and 0.2% DSS-treated C57BL/6 mice to antibiotic-treated mice [43]. The recipient mice were denoted FM*_NC_* and FM*_DSS_* mice, respectively. Two weeks after the transplantation, the FM*_DSS_* and FM*_NC_* mice developed a gut microbiota more similar to those of their donor (Figure S7). Moreover, the FM_DSS_ mice showed significantly higher blood glucose level, lower insulin secretion, more beta cell destruction and more signs of pancreatitis compared with the FM*_NC_* mice, even though all the recipient mice were provided pure water during the trial (Figure 2, Figure S2). This result demonstrates that the gut microbiota treated by low dose DSS is sufficient to induce IDD in mice. Thus, this chemically induced IDD in mice was causally mediated by the gut microbiota which was disrupted by the low dose DSS treatment.

**Figure 2.**
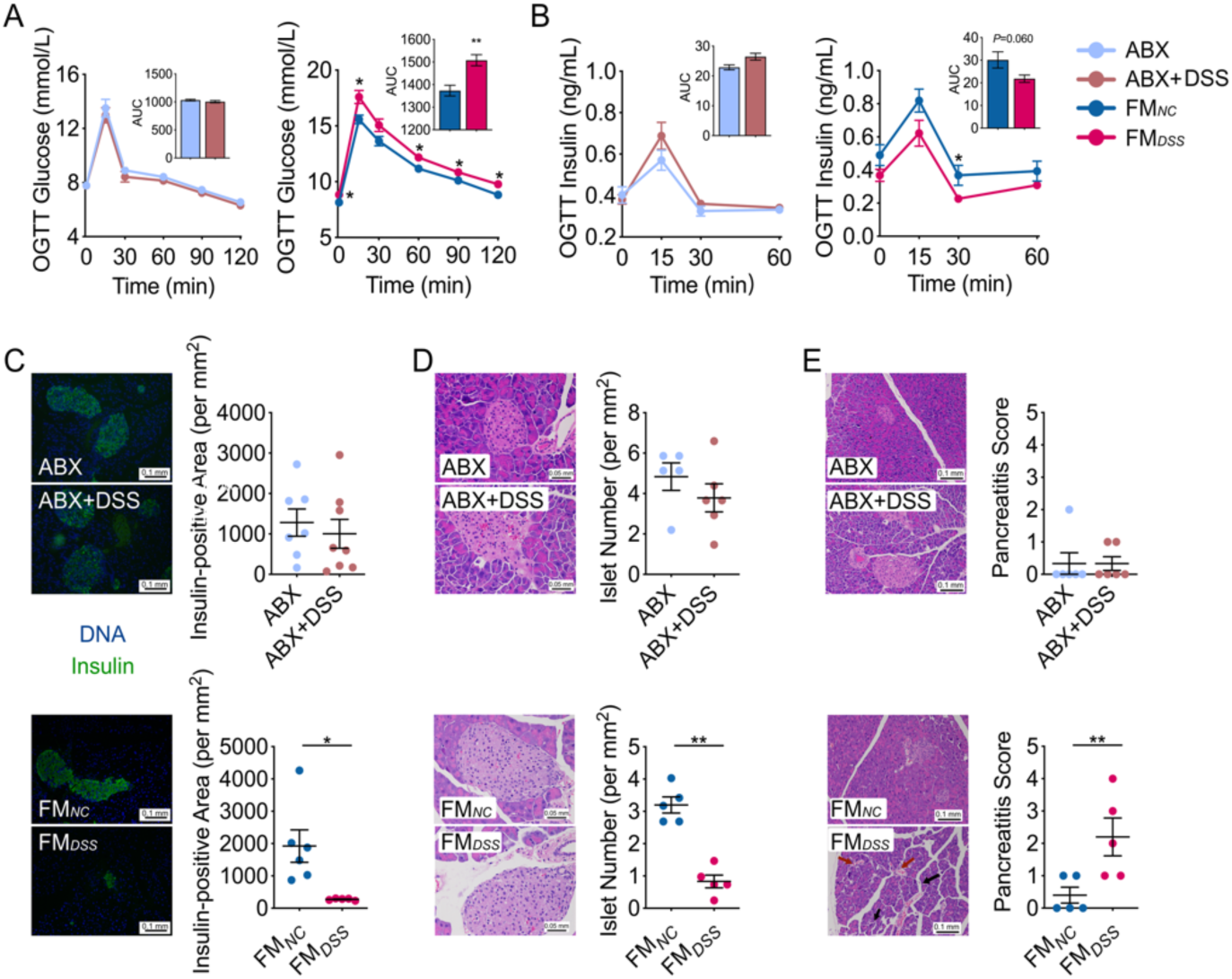
The gut microbiota disrupted by 0.2% DSS was both necessary and sufficient for inducing IDD. **(A)** Blood glucose and **(B)** insulin levels with the AUC in the OGTT of ABX, ABX+DSS, FM*_DSS_* and FM*_NC_* mice. For **(A)**, ABX group, n=10; ABX+DSS group, n=8; FM*_NC_* group, n=8; FM*_DSS_* group, n=8. For **(B)**, ABX group, n=9; ABX+DSS group, n=7; FM*_NC_* group, n=6; FM*_DSS_* group, n=6. **(C)** Histological sections of pancreatic tissue (200×, scale bar = 0.1 mm) from ABX, ABX+DSS, FM*_DSS_* and FM*_NC_* mice stained for insulin (green) and DNA (blue), and the insulin-positive area was quantified (at bottom). ABX group, n=7; ABX+DSS group, n=8; FM*_NC_* group, n=6; FM*_DSS_* group, n=5. **(D)** The number of islets per mm^2^ of pancreatic tissue was calculated based on H&E-stained histological sections of pancreatic tissue (400×, scale bar = 0.05 mm) from ABX, ABX+DSS, FM*_DSS_* and FM*_NC_* mice. ABX group, n=5; ABX+DSS group, n=6; FM*_NC_* group, n=5; FM*_DSS_* group, n=5. **(E)** The histologic pancreatitis score was evaluated by edema, inflammation, and vacuolization in H&E-stained histological sections of pancreatic tissue (200×, scale bar = 0.1 mm) from ABX, ABX+DSS, FM*_DSS_* and FM*_NC_* mice. Black arrows, scattered structure; Red arrows, increased peri-islet neovascularization. ABX group, n=6; ABX+DSS group, n=6; FM*_NC_* group, n=5; FM*_DSS_* group, n=5. The data in **(A)-(E)** are shown as the mean ± SEM, and Student’s t-test (two-tailed) was used to analyze the following pairs of groups: ABX vs. ABX+DSS, or FM*_NC_* vs. FM*_DSS_*. \**P* <0.05 and \*\**P* <0.01. ABX, antibiotic cocktail-treated mice provided pure water; ABX+DSS, antibiotic cocktail-treated mice treated with 0.2% (w/v) DSS added to drinking water; FM*_NC_*, antibiotic cocktail-treated mice inoculated with the fecal microbiota from mice in the NC group and provided pure water; FM*_DSS_*, antibiotic cocktail-treated mice inoculated with the fecal microbiota from mice in the DSS group and provided pure water.

### Translocation of candidate gut pathobionts to the pancreas promoted local inflammation

To identify key members of the gut microbiota enriched by the low dose DSS which may causally contribute to IDD development, we first identified ASVs in the gut microbiota which showed significant differences between the 0.2% DSS-treated and control mice by sparse partial least squares discriminant analysis (sPLS-DA) (Figure 3A and Figure S8). Notable changes were observed in the family Muribaculaceae (phylum Bacteroidetes), a major but understudied mouse taxon [44]. Specifically, 4 ASVs in this family were enriched, while 6 ASVs were depleted (Figure 3A). On the other hand, ASVs associated with potential butyrate producers, such as 2 ASVs in the genus *Eubacterium*, were decreased (Figure 3A). Butyrate content was significantly lower in DSS-treated than in control mice (Figure 3B). The butyrate content was also significantly lower in FM*_DSS_* mice compared with that in FM*_NC_* mice (Figure 3B). Reduced diversity of butyrate-producing bacteria and lower gut butyrate content have been linked with overgrowth of pathobionts [45]. We hypothesized that the 4 ASVs in the family Muribaculaceae enriched by low dose DSS may represent candidate pathobionts for IDD development.

**Figure 3.**
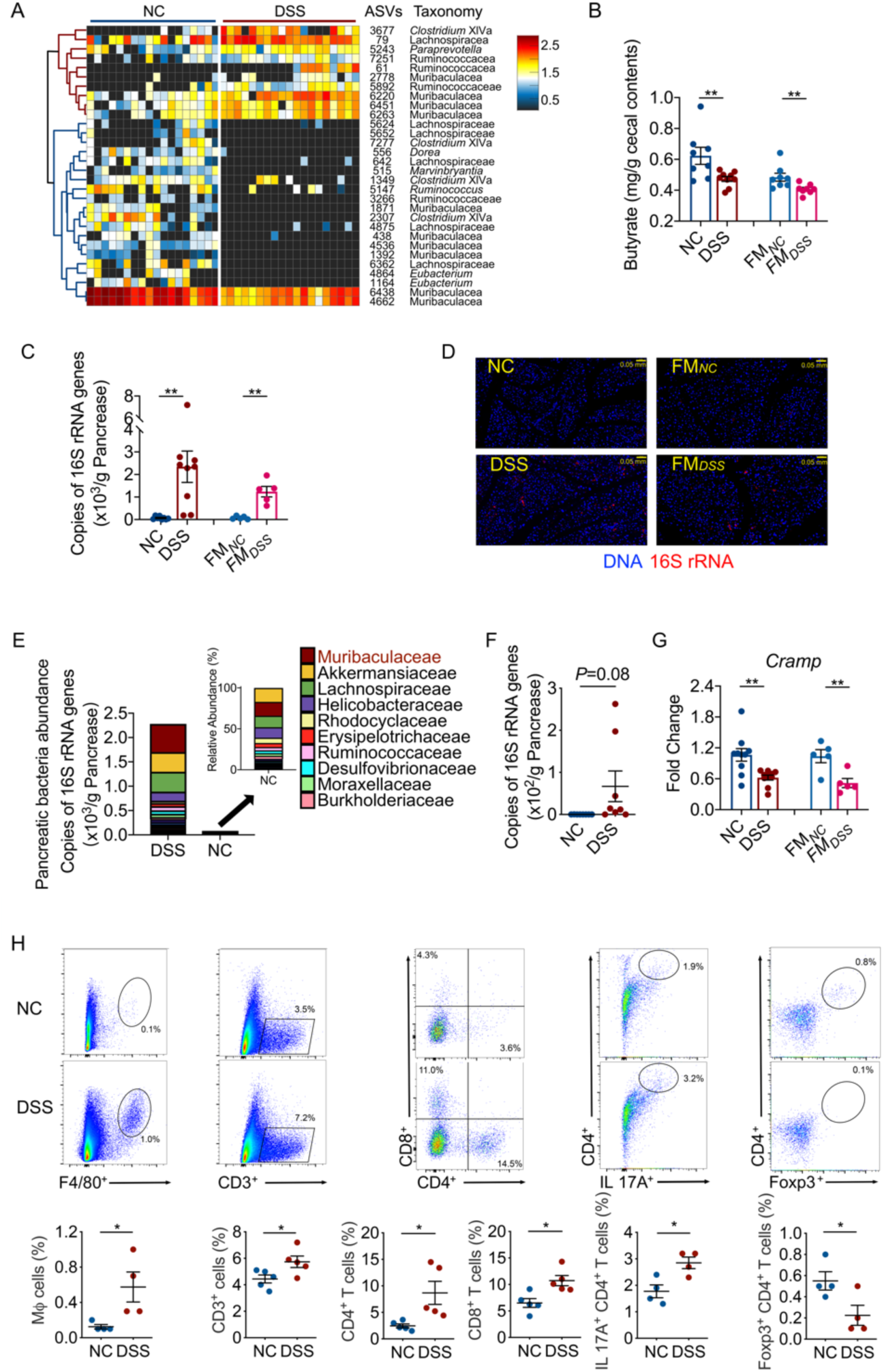
An increase in bacterial load and the impairment of immune tolerance in the pancreas. **(A)** Thirty ASVs that were significantly altered after 3 months of 0.2% DSS treatment, as identified using sPLS-DA models. The cluster tree on the left shows associations between these ASVs, as determined by the Spearman correlation coefficient based on their relative abundances among all the samples. The heat map shows the relative abundance (log_10_ transformed) of each ASV in a sample from an individual mouse. NC group, n=17; DSS group, n=18. **(B)** The butyrate content in the cecum of mice. NC, n=8; DSS, n=9; FM*_NC_* n=8; FM*_DSS_*, n=8. The bacterial load in the pancreas as measured by **(C)** real-time qPCR and **(D)** fluorescence in situ hybridization (FISH) of 16S rRNA. NC, n=8; DSS, n=9; FM*_NC_*, n=5; FM*_DSS_*, n=5. **(E)** Distribution of family-level phylotypes in the pancreas of NC and DSS-treated mice. The names of the top 10 most abundant families in the DSS group are shown. The abundance was determined by the relative abundances of these families with the total bacterial load in the pancreas detected with qPCR of 16S rRNA genes. NC, n=9; DSS, n=8. **(F)** The load of the 4 Muribaculaceae ASVs in the pancreas. The load was determined by multiplying the relative abundances of these ASVs with the total bacterial load in the pancreas detected with qPCR of 16S rRNA genes. NC, n=9; DSS, n=8. **(G)** *Cramp* expression in the pancreas of mice. NC, n=10; DSS, n=10; ABX, n=6; ABX+DSS, n=6; FM*_NC_*, n=5; FM*_DSS_*, n=5. **(H)** Flow cytometry evaluation of leukocyte subtypes in the pancreas of NC and DSS-treated mice. Data are presented as the frequency of gated cells (from left to right: F4/80^+^ macrophages, CD3^+^ cells, CD8^+^ T cells, CD8^+^ IFNγ^+^ cells, CD4^+^ T cells, IL-17A^+^ CD4^+^ cells, and Foxp3^+^ CD4^+^ cells) among the CD45^+^ population of cells per mouse. NC group, n=4-5; DSS group, n=4-5. The data are shown as the mean ± SEM, and Student’s t-test (two-tailed) was used to analyze differences between the NC and DSS groups or the FM*_NC_* and FM*_DSS_* groups. \**P* <0.05 and \*\**P* <0.01.

Translocation of gut pathobionts to pancreas has bene linked with a few disease conditions [46]. We hypothesized that the candidate pathobionts enriched in low dose DSS disrupted gut microbiota may translocate to pancreas and induce IDD. Indeed, qPCR and fluorescence in situ hybridization (FISH) targeting the 16S rRNA gene showed that the bacterial load was significantly higher (by at least one order of magnitude) in the pancreas of the low dose DSS-treated and FM*_DSS_* mice than in that of their respective controls (Figures 3C and 3D, Figure S9 and S10). Absolute abundance by multiplying the relative abundance from sequencing the V3-V4 region of the 16S rRNA gene of the microbiota with qPCR derived total copy number of 16S rRNA genes in the pancreas showed that the most abundant bacteria in the pancreas of control mice belonged to the family Akkermansiaceae, but the microbiota in the pancreas of low dose DSS-treated mice was dominated by the family Muribaculaceae (Figure 3E). Notably, the 4 ASVs in Muribaculaceae, which were significantly enriched in the gut by 0.2% DSS treatment (Figure 3A), also showed a trend of being enriched in the pancreas of the low dose DSS-treated mice (Figure 3F). Other ASVs in the same family showed no difference in pancreas between the low dose DSS-treated and control mice. The bacterial load was not increased in the liver of the low dose DSS-treated mice (Figure S11), suggesting that the gut microbiota specifically translocated to the pancreas. Reduced production of cathelicidin-related antimicrobial peptide (CRAMP) by pancreatic beta cells has been linked with the development of autoimmunity in the pancreas and T1D [16]. We found that the expression of the *Cramp* gene in the pancreas of 0.2% DSS-treated mice and FM*_DSS_* mice was significantly lower than that in the pancreas of their respective controls (Figure 3G). The reduced antimicrobial peptide production may have allowed bacterial overgrowth in pancreas. Thus, the low dose DSS enriched ASVs in the gut microbiota translocated to and overgrew in pancreas.

We then examined immune responses of pancreas to the bacterial translocation and overgrowth. Infiltration of macrophages and total T cells were higher in the pancreas of the low dose DSS-treated mice than control (Figure 3H). T cell differentiation toward CD4^+^ and CD8^+^ T cells and T_H_17 cells were also higher in low dose DSS-treated mice than control, which may contribute to more inflammatory tissue damage [47]. Moreover, the low dose DSS-treated mice had a decreased population of regulatory T (T_reg_) cells, which suppress the pathological immune response [48], suggesting a continued inflammatory pancreatic milieu (Figure 3H). These immune responses were also observed in FM_DSS_ mice but not in FM*_NC_* mice (Figure S12). ABX+DSS and ABX mice exhibited no detectable pancreatic bacterial loads via qPCR and FISH and no differences in immune responses (Figure S13). Taken together, our data supports the hypothesis that the downregulation of the pancreatic antimicrobial peptide CRAMP may allow more bacteria to translocate from the gut to the pancreas and overgrew there, which may lead to immune tolerance disruption and IDD development.

### A pathobiont isolate from the family Muribaculaceae induced IDD in germfree mice

Based on above findings, we hypothesized that specific bacteria represented by the 4 ASVs in the family Muribaculaceae, which were enriched in both the gut and pancreas, might represent the key mediators of low dose DSS-induced IDD. To further examine the causal role of the bacteria in this family which were associated with low dose DSS-induced IDD, we first aimed to obtain their pure cultures from fresh feces of the low dose DSS-treated mice. Although bacteria in Muribaculaceae are dominant in the mouse gut microbiota, they had not been cultured until recently [44]. After sequence-guided screening of 13,079 colonies isolated using 46 different types of media (Table S1), we obtained a pure culture (named strain MF 13079) and its finished genomic sequence (Figure 4A and 4B). The genome of the strain MF 13079 had two different 16S rRNA genes, each of which had 100% identity in the V3-V4 region with ASV 6263 and ASV 6451 respectively (Figure S14). These two ASVs were members of the family Muribaculaceae, which were among the 4 ASVs enriched in both the gut and pancreas of low dose DSS-treated mice (Figure 3A). The nearest neighbor of MF 13079 in GenBank is *Duncaniella freteri* strain TLL-A3 but with only 93.7% homology in 16S rRNA gene sequence (Figure 4C). The average nucleotide identity (ANI) values between the genome of strain MF 13079 and all 10 publicly available genomes of Muribaculaceae in GenBank ranged from 73.3% to 75.1%. Thus, MF 13079 may represent a new genus in Muribaculaceae [49]. Moreover, we found the genes for protecting against oxidative damage (COG0605 superoxide reductase, COG0753 catalase, COG0492 thioredoxin reductase, COG0450/COG2077/COG1225 peroxiredoxin) and genes for adhesion and immune evasion (COG0133/COG0159 tryptophan synthase) in the genome of MF 13079 (Table S2). Such genes may facilitate the overgrowth of this gut bacterium in the pancreas and qualify it as a pathobiont [50].

**Figure 4.**
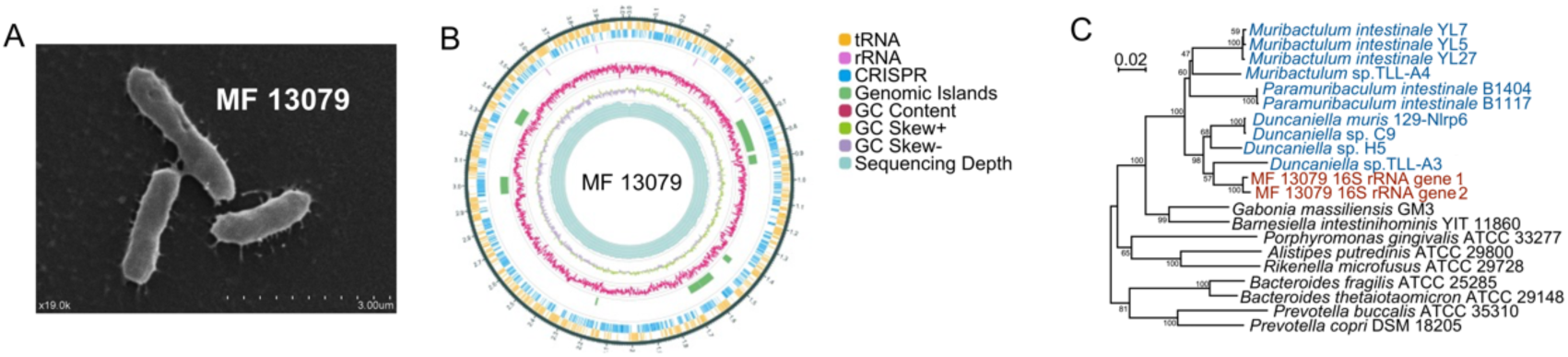
A gut-derived Muribaculaceae isolate enriched by 0.2% DSS treatment. **(A)** A scanning electron microscopy image of a pure isolate from the Muribaculaceae family (named strain MF 13079) that was enriched in the gut of DSS-treated mice. **(B)** Genome atlas of the MF13079 strain of Muribaculaceae. From outer to inner: circle 1. tRNAs; circle 2. rRNAs; circle 3. CRISPR sites; circle 4. mean centered GC content; circle 5. Sequencing depth. **(C)** A phylogenetic tree constructed with the 16 rRNA gene of MF 13079 (red), other Muribaculaceae bacteria (blue) and typical bacterial strains in Bacteroidetes.

To test if MF 13079 by itself can translocate from the gut to pancreas and induce IDD, we inoculated only MF 13079 into germ-free adult C57BL/6 mice (Muri) by oral gavage. A mouse-derived strain, *Akkermansia muciniphila* 139, was inoculated into germ-free mice as control (Akk) [51]. Butyrate was not detected in either group (Figure 5A), and the pancreatic *Cramp* gene expression level showed a trend of being lower in Muri group (Figure 5B). MF 13079 attained a significantly higher load in the pancreas of Muri mice than *A. muciniphila* 139 in that of Akk mice (Figure 5C), suggesting that the capacity of MF 13079 to translocate to the pancreas is much higher than that of *A*. *muciniphila* 139. The Muri group, but not the Akk group, showed inflammation in the pancreas and development of IDD (Figure 5D-5F). Thus, MF 13079 alone can translocate from the gut to the pancreas, overgrew there and induce IDD.

**Figure 5.**
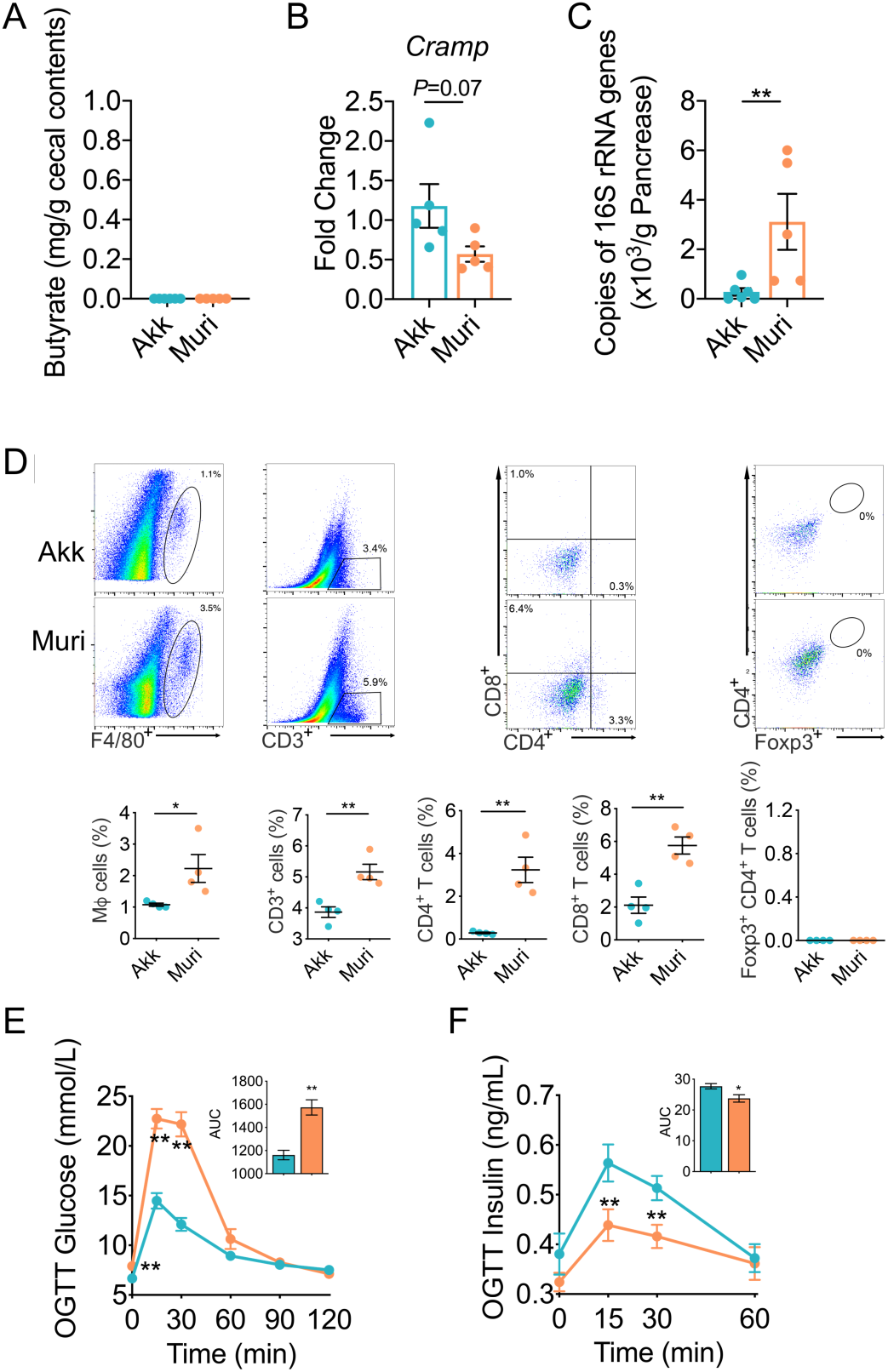
MF13079 induced IDD by translocating to the pancreas in germ-free mice. **(A)** The butyrate content in the cecum of mice in the Muri and Akk groups. Akk, n=12; Muri, n=11. **(B)** *Cramp* expression in the pancreas of mice in the Muri and Akk groups. Akk, n=6; Muri, n=5. **(C)** The MF 13079 load in the pancreas of mice in the Muri and Akk groups assessed via qPCR of the 16S rRNA gene of each strain. Akk, n=6; Muri, n=5. **(D)** Flow cytometry evaluation of leukocyte subtypes in the pancreas of mice in the Muri and Akk groups. Data are presented as the frequency of gated cells (left to right: F4/80^+^ macrophages, CD3^+^ cells, CD8^+^ T cells, CD4^+^ T cells, and Foxp3^+^ CD4^+^ cells) among the CD45^+^ population of cells per mouse. Akk, n=5; Muri, n=5. Blood glucose **(E)** and insulin **(F)** levels of the mice in Muri and Akk groups as measured by OGTT (the AUC is shown in the inset graph). For **(E)**, Akk, n=12; Muri, n=11. For **(F)**, Akk, n=10; Muri, n=10. Muri, germ-free mice inoculated with the MF 13079 strain and provided pure water; Akk, germ-free mice inoculated with the *Akkermansia muciniphila* strain 139 and provided pure water. In **(A)-(F)**, the data are shown as the mean ± SEM, and Student’s t-test (two-tailed) was used to analyze the differences between the Muri and Akk groups. \**P* <0.05 and \*\**P* <0.01.

To test whether MF 13079 can translocate from the gut to the pancreas and induce IDD when in competition with a complex normal gut microbiota and in a host with normal immunity, we transplanted cultured MF 13079 together with feces from control mice by oral gavage into antibiotic-treated mice (FM*_NC+Muri_*). Antibiotic-treated mice inoculated with feces from control mice were denoted as FM*_NC_*. We found that FM*_NC+Muri_* mice had significantly lower cecum butyrate levels and pancreatic *Cramp* expression than FM*_NC_* mice (Figure 6A and 6B). We obtained the absolute abundance of MF 13079 by multiplying the relative abundances of ASVs 6263 and 6451 with the total bacterial load in the pancreas detected with qPCR of 16S rRNA genes. We found that the load of MF 13079 in the pancreas of FM*_NC+Muri_* mice was three orders of magnitude higher than that in the pancreas of FM*_NC_* mice (Figure 6C). Concomitantly, the FM*_NC+Muri_* mice showed an inflammatory pancreatic milieu and increased blood glucose and decreased insulin in the OGTT (Figure 6D-6F). Thus, when transplanted together with the fecal microbiota from healthy mice, MF 13079 still translocated to pancreas and induced IDD in antibiotic-treated mice.

**Figure 6.**
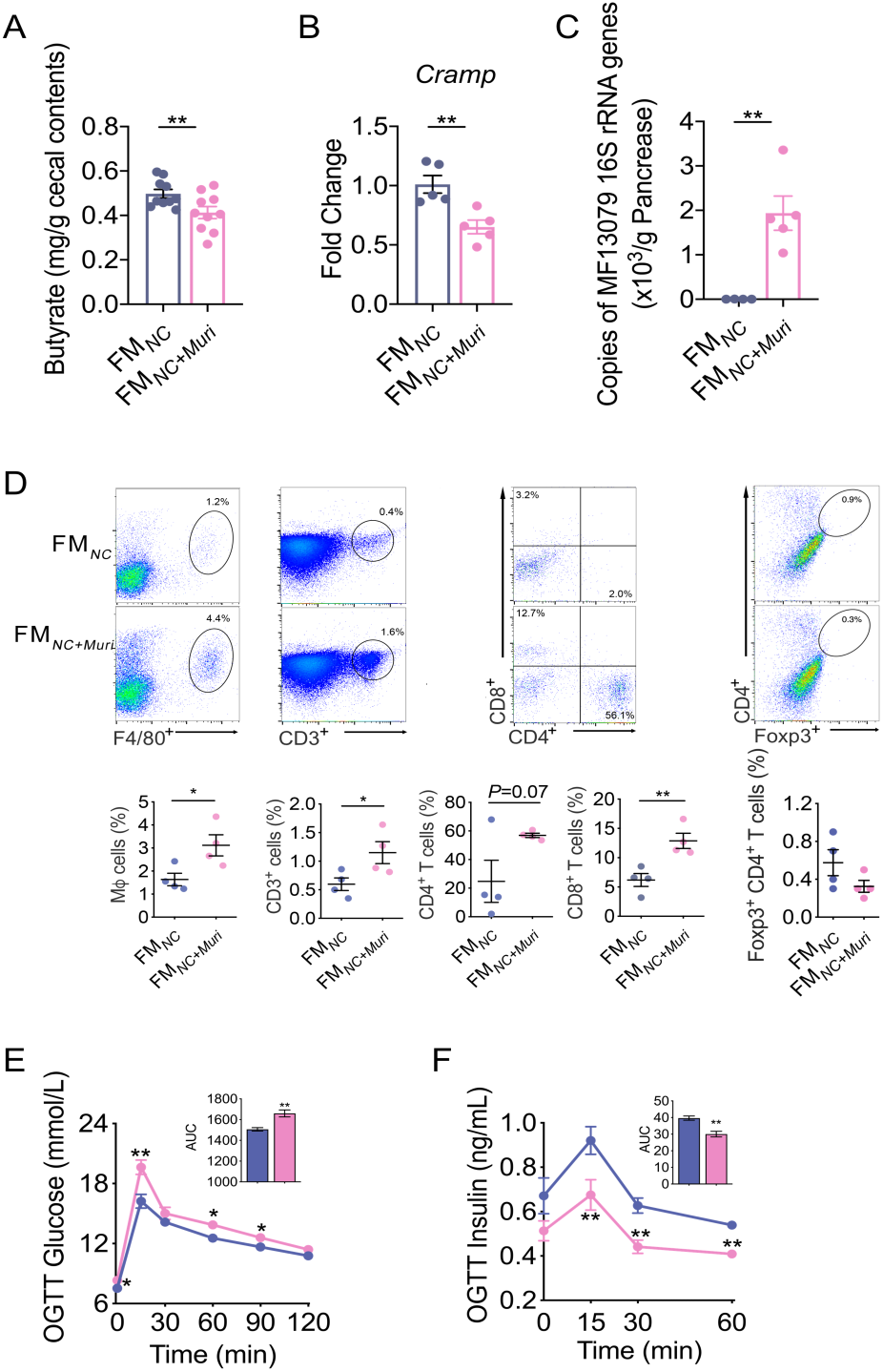
MF 13079 translocated from the gut to the pancreas and induce IDD when in competition with a complex gut microbiota. **(A)** The butyrate content in the cecum of FM*_NC_* and FM*_NC+Muri_* mice. FM*_NC_*, n=10; FM*_NC+Muri_*, n=10. **(B)** *Cramp* expression in the pancreas of FM*_NC_* and FM*_NC+Muri_* mice. FM*_NC_*, n=5; FM*_NC+Muri_*, n=5. **(C)** The amount of the MF 13079 in the pancreas of FM*_NC+Muri_* and FM*_NC_* mice assessed with the method described in Figure 3F. FM*_NC_*, n=4; FM*_NC+Muri_*, n=5. **(D)** Flow cytometry evaluation of leukocyte subtypes in the pancreas of FM*_NC+Muri_* and FM*_NC_* mice. Data are presented as the frequency of gated cells (left to right: F4/80^+^ macrophages, CD3^+^ cells, CD8^+^ T cells, CD4^+^ T cells, and Foxp3^+^ CD4^+^ cells) among the CD45^+^ population of cells per mouse. FM*_NC_*, n=4; FM*_NC+Muri_*, n=4. Blood glucose **(E)** and insulin **(F)** levels of FM*_NC+Muri_* and FM*_NC_* mice as measured by an OGTT (the AUC is shown in the inset graph). For **(E)**, FM*_NC_*, n=10; FM*_NC+Muri_*, n=10. For **(F)**, FM*_NC_*, n=8; FM*_NC+Muri_*, n=10. FM*_DSS_*, antibiotic cocktail-treated mice inoculated with the fecal microbiota from mice in the DSS group and provided pure water; FM*_NC+Muri_*, antibiotic cocktail-treated mice inoculated with a mixture of the fecal microbiota of the NC group and the MF 13079 strain and provided pure water. In **(A)-(F)**, the data are shown as the mean ± SEM, and Student’s t-test (two-tailed) was used to analyze the differences between the FM*_NC_* and FM*_NC+Muri_* groups. \**P* <0.05 and \*\**P* <0.01.

As recipient mice had normal gut barrier function, we tested other possible routes for translocation of gut pathobionts to pancreas. Notably, MF 13079 has a complete flagellar assembly pathway in its genome based on KEGG database querying, and a flagellar structure was observed with scanning electron microscopy (Figure 4A, Figure S15 and Table S2). We tested bacterial motility by an agar-based motility assay, and MF 13079 demonstrated notable motility in vitro (Figure S15). It provides a means by which MF 13079 might translocate from the gut to the pancreas via the pancreatic duct. Interestingly, we also detected a much higher load of MF 13079 in the mesenteric lymph nodes of Muri mice than *A. muciniphila* 139 in those of Akk mice (Figure S16), which might represent another pathway for MF 13079 to gain access to the pancreas via lymphatic drainage routes [52, 53].

## Discussion

Our study demonstrated that novel pathobionts enriched in a low DSS-disrupted gut microbiota can translocate to pancreas and induce IDD in mice. Such pathobionts are both necessary and sufficient to induce IDD. Genetic predisposition to IDD here was not required from the host side as the mice we used were with no known relevant genetic defects. More importantly, a single pathobiont MF 13079 isolated from the family Muribaculaceae was sufficient to induce IDD when mono-associated with wildtype germfree mice. The fact that IDD was induced when MF 13079 was mixed with normal gut microbiota and transplanted to antibiotic treated wildtype mice which have well-developed immune system shows that predominant population level of this pathobiont in the gut is sufficient to IDD development. Thus, this pathobiont has been identified as an etiological factor for inducing IDD as what we did fulfilled Koch’s postulates which were established for identifying the causal agent for an infectious disease. This IDD-inducing capacity may be a redundant function of many gut pathobionts in the same mouse, however, translocation of one such pathobiont to pancreas seems to be enough for inducing IDD and host genetic defect is not required.

Studies in humans have shown that acute or chronic inflammation of the entire pancreas may drive beta-cell death and lead to IDD [54]. Previous studies showed that decreased gene expression of pancreatic CRAMP stimulated the inflammatory activities of intrapancreatic immune cells and led to destruction of beta cells [16]. Our study showed that enrichment and overgrowth of pathobionts in the pancreas is necessary for CRAMP-induced stimulation of local immune cells because antibiotic removal of the gut microbiota resulted in pancreatic microbiome depletion and abolished DSS-induced IDD.

Studies showed that reduced butyrate production in the gut may drive the decreased gene expression of pancreatic CRAMP [?]. This might be a mechanism for induction of pancreatic inflammation and IDD in our low DSS model. The low dose DSS treated mice had depletion of butyrate producers and reduced butyrate production in their gut and decreased gene expression of pancreatic CRAMP. This warrants further studies.

## Conclusion

Our study shows that toxic chemicals such as DSS may disrupt the gut microbiota and lead to serious health consequences even at a low dosage which does not cause acute local responses in the gut. The overgrowth in the gut and translocation to the pancreas of pathobionts may induce inflammation in the pancreas, destruct beta cells and lead to IDD. This study inspires us to search for similar pathobionts in patients with IDD. Suppression of the overgrowth of pathobionts in the gut and prevention of their translocation to the pancreas via restoration of a normal structure of the gut microbiota may become a promising ecological approach for managing IDD.

## List of abbreviation

IDD: Insulin-dependent diabetes
T1D: type 1 diabetes
SCFAs: short-chain fatty acids
DSS: dextran sulfate sodium
OGTT: oral glucose tolerance test
ITT: insulin tolerance test
PCoA: principal coordinate analysis
ASVs: amplicon sequence variants
PERMANOVA: permutational multivariate analysis of variance
sPLS-DA: sparse partial least squares discriminant analysis
FISH: fluorescence in situ hybridization
CRAMP: cathelicidin-related antimicrobial peptide

## Declarations

### Ethics approval and consent to participate

All experimental procedures for specific pathogen free (SPF) mice were approved by the Institutional Animal Care and Use Committee (IACUC) of the School of Life Sciences and Biotechnology of Shanghai Jiao Tong University (no. 2014005). The procedures for the germ-free mice experiments were approved by the Institutional Animal Care and Use Committee of SLAC Inc (no. 20190201001).

### Consent for publication

Not applicable

### Availability of data and materials

The data that support the findings of this study are available from the corresponding author upon request. The 16S rRNA gene sequences of Muribaculaceae strain MF13079 has been submitted to the GenBank with the accession No. MT921595 and MT921596. The whole-genome sequence of Muribaculaceae strain MF13079 has been submitted to the NCBI under accession No. PRJNA576346.The raw Illumine sequence data generated in this study are available in the sequence read archive (SRA) at NCBI under accession No. SRP278697.

### Founding

This work was supported by grants from the National Natural Science Foundation of China (31922003 and 81871091) and the National Key Research and Development Project (2019YFA0905600).

## Acknowledgements

Not applicable

## Author’s contributions

C. Z. and L. Z. conceived, planned and supervised this study. X. Y. carried out the animal experiments and most experiments in lab. X. X. and Z. R. isolated the *Akkermansia muciniphila* strain 139 and provided the protocol of *A. muciniphila* strain 139 related animal experiment. X. Y. and J. N. performed the flow cytometry evaluation of subtypes of leukocytes in the pancreas. G. W. and Z. W. provided help of genome sequencing data analysis. C. Z. and X. Y. analyzed the data and generated the figures and tables. C. Z., L. Z., H. Y., J. Y., X. Y. and G. W. prepared the manuscript. All authors approved the manuscript.

## Supplementary Files

1. File name: Supplementary Figures File Format: PDF

Discerption: Supplementary Figure S1 to S16

Fig. S1| Fasting C-Peptide in the mice with and without 0.2% DSS.

Fig. S2| Serum amylase (AMS) and serum lipase levels in each group.

Fig. S3| 0.2% DSS increased food and water intake but did not change energy intake of mice.

Fig. S4| 0.2% DSS did not change the lipid accumulation in mice.

Fig. S5| 0.2% DSS did not impair gut barrier integrity and induce inflammation.

Fig. S6| The gut microbiota had been depleted by more than 99% in the mice with a cocktail of antibiotics for 5 weeks.

Fig. S7| The recipient mice developed a gut microbiota more similar to their donor mice.

Fig. S8| Optimal classification performance of the sPLS-DA model of the gut microbial structure in NC and DSS mice.

Fig. S9| Fluorescence in situ hybridization (FISH) of 16S rRNA in the pancreas of antibiotic treated mice.

Fig. S10| Fluorescence in situ hybridization (FISH) of 16S rRNA in the pancreas in fecal microbiota transplanted mice.

Fig. S11| The bacterial load in the liver as measured by real-time qPCR of 16S rRNA gene.

Fig. S12| Flow cytometry evaluation of subtypes of leukocytes in the pancreas of FM*_NC_* and FM*_DSS_* mice.

Fig. S13| 0.2% DSS did not enrich bacteria in pacreas and disrupt the immune tolerance in antibiotic treated mice.

Fig. S14| Identification of Muribaculaceae strain MF 13079.

Fig. S15| Functional annotation of flagella-related genes and motility test of MF13079.

Fig. S16| Bacteria load in mesenteric lymph nodes (MLN) in Akk and Muri mice.

2. File name: Tables S1

File Format: PDF

Title and Discerption: Culture medium used for isolation of Muribaculeae

3. File name: Tables S2

File Format: Excel

Title and Discerption: COG in the genome of MF 13079

