## Supplementary Figure 1-16 for "Pathobionts from chemically disrupted gut microbiota induce insulin-dependent diabetes in mice"

#### Summary

1. File name: Supplementary Figures

   File Format: PDF

   Discerption: Supplementary Figure S1 to S16

   Fig. S1| Fasting C-Peptide in the mice with and without 0.2% DSS.

   Fig. S2| Serum amylase (AMS) and serum lipase levels in each group.

   Fig. S3| 0.2% DSS increased food and water intake but did not change energy intake of mice.

   Fig. S4| 0.2% DSS did not change the lipid accumulation in mice.

   Fig. S5| 0.2% DSS did not impair gut barrier integrity and induce inflammation.

   Fig. S6| The gut microbiota had been depleted by more than 99% in the mice with a cocktail of antibiotics for 5 weeks.

   Fig. S7| The recipient mice developed a gut microbiota more similar to their donor mice.

   Fig. S8| Optimal classification performance of the sPLS-DA model of the gut microbial structure in NC and DSS mice.

   Fig. S9| Fluorescence in situ hybridization (FISH) of 16S rRNA in the pancreas of antibiotic treated mice.

   Fig. S10| Fluorescence in situ hybridization (FISH) of 16S rRNA in the pancreas in fecal microbiota transplanted mice.

   Fig. S11| The bacterial load in the liver as measured by real-time qPCR of 16S rRNA gene.

Fig. S12| Flow cytometry evaluation of subtypes of leukocytes in the pancreas of $FM_{NC}$ and $FM_{DSS}$ mice.

Fig. S13| 0.2% DSS did not enrich bacteria in pacreas and disrupt the immune tolerance in antibiotic treated mice.

Fig. S14| Identification of Muribaculaceae strain MF 13079.

Fig. S15| Functional annotation of flagella-related genes and motility test of MF13079.

Fig. S16| Bacteria load in mesenteric lymph nodes (MLN) in Akk and Muri mice.

2. File name: Tables S1

   File Format: PDF

   Title and Discerption: Culture medium used for isolation of Muribaculeae

3. File name: Tables S2

   File Format: Excel

   Title and Discerption: COG in the genome of MF 13079

### Figures

##### Figure S1

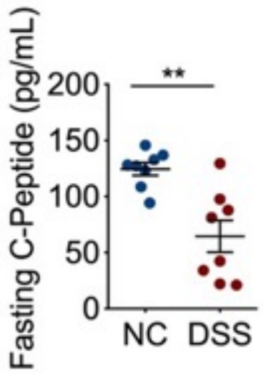


**Fig. S1 Fasting C-Peptide in the mice with and without 0.2% DSS.** NC group, n=8; DSS group, n=8. NC, the control mice treated with pure drinking water; DSS, the mice were treated with 0.2% (w./v.) DSS added in drinking water. The data in were shown as mean ± s.e.m., and Student's t-test (two-tailed) was used to analyze differences between groups. ***P* <0.01.

**Figure S2**

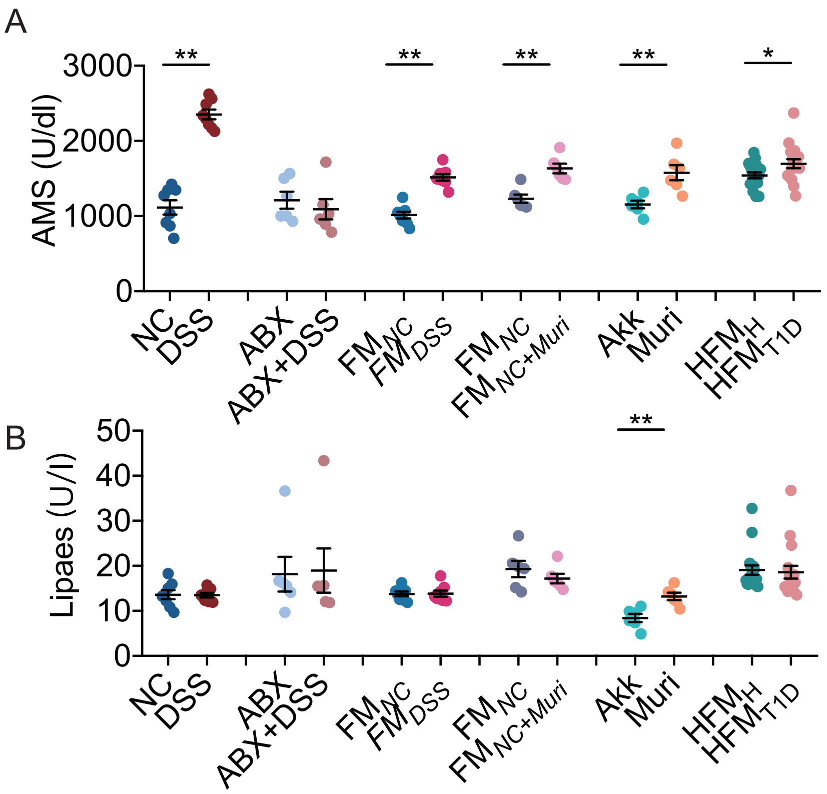


**Fig. S2** Serum amylase (AMS) **(A)** and serum lipase **(B)** levels in each group. NC group, n=8; DSS group, n=8; ABX group, n=6; ABX+DSS group, n=6; FM$_{NC}$ group, n=8; FM$_{DSS}$ group, n=8; FM$_{NC}$ group, n=6; FM$_{NC+Muri}$ group, n=6; Akk group, n=6; Muri group, n=6; HFM$_{H}$ group, n=18; HFM$_{T1D}$ group, n=17. NC, the control mice treated with pure drinking water; DSS, the mice were treated with 0.2% (w./v.) DSS added in drinking water; ABX, the antibiotic-cocktail treated mice with pure water; ABX+DSS, the antibiotic-cocktail treated mice with 0.2% (w./v.) DSS added in drinking water; FM$_{NC}$, the antibiotic-cocktail treated mice inoculated with the fecal microbiota from the mice of NC group, and had pure water; FM$_{DSS}$, the antibiotic-cocktail treated mice inoculated with the fecal microbiota from the mice of DSS group, and had pure water; FM$_{NC+Muri}$, the antibiotic-cocktail treated mice inoculated with mixture of the fecal microbiota of NC group and MF 13079 strain, and had pure water; Muri, the germ-free mice inoculated with the MF 13079 strain, and had pure water; Akk, the germ-free mice inoculated with the Akkermansia muciniphila strain 139, and had pure water. Data were shown as mean ± s.e.m., and Student's t-test (two-tailed) was used to analyze differences between NC vs. DSS, ABX vs. ABX+DSS, FM$_{NC}$ vs. FM$_{DSS}$, FM$_{NC}$ vs. FM$_{DSS+Muri}$, Akk vs. Muri, HFM$_{H}$ vs. HFM$_{T1D}$. *$P$ <0.05 and **$P$ <0.01.

**Figure S3**

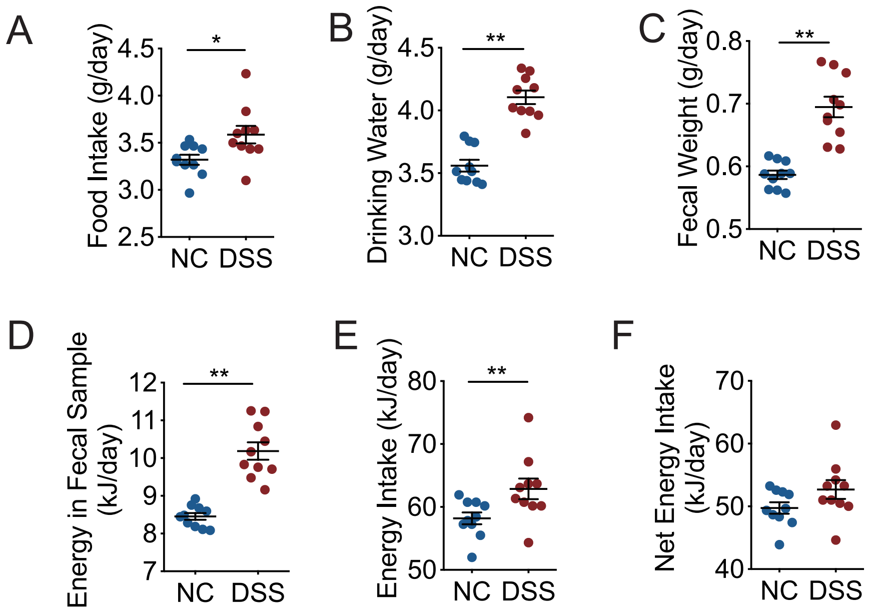


**Fig. S3 0.2% DSS increased food and water intake but did not change energy intake of mice.** (**A**) Food intake, **(B)** drinking water, **(C)** fecal weight, **(D)** fecal energy content, **(E)**energy intake and **(F)** net energy intake of NC and DSS mice. NC group, n=8; DSS group, n=9. NC, the control mice treated with pure drinking water; DSS, the mice were treated with 0.2% (w./v.) DSS added in drinking water. The data in **(A)-(G)** were shown as mean ± s.e.m., and Student's t-test (two-tailed) was used to analyze differences between groups. **P* <0.05 and ***P* <0.01.

**Figure S4**

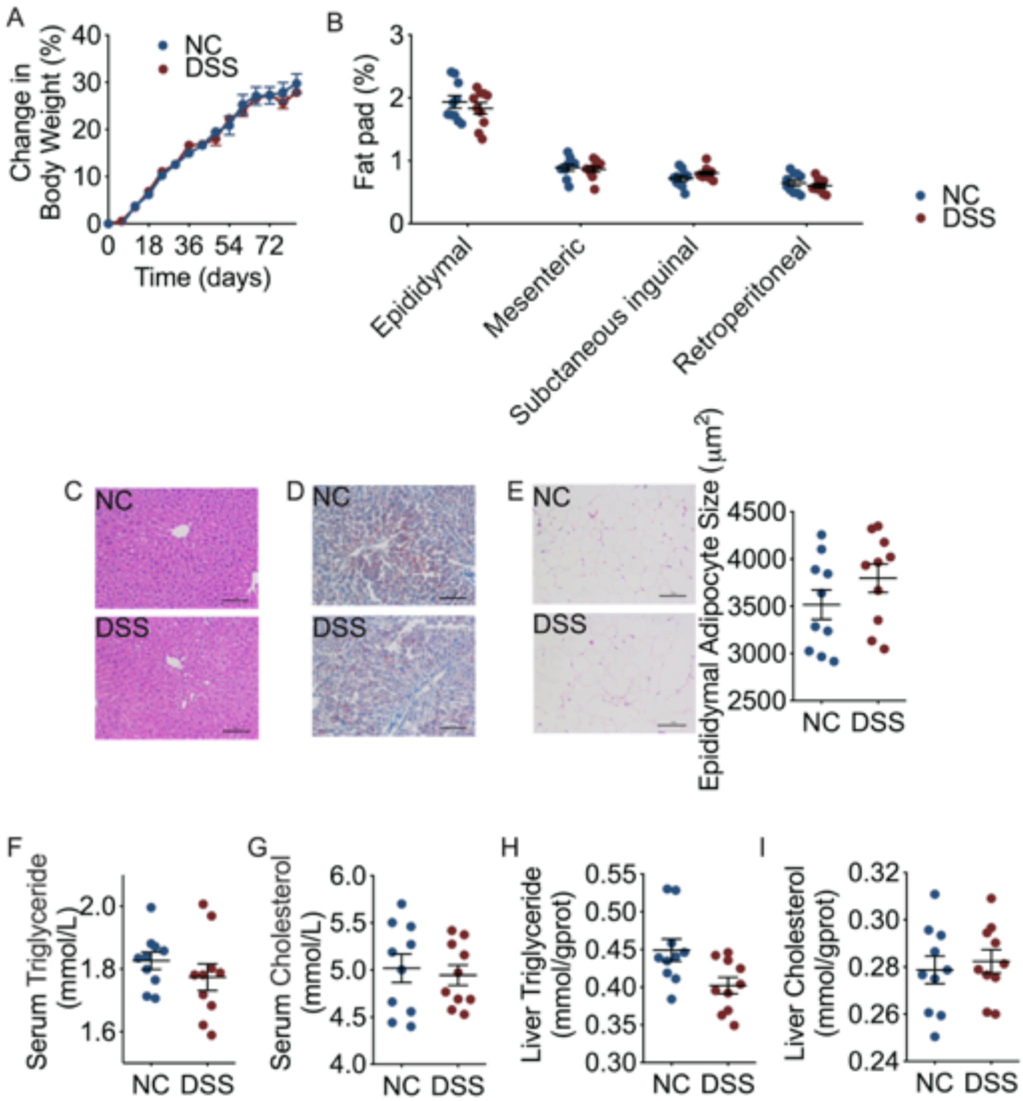


**Fig. S4 0.2% DSS did not change the lipid accumulation in mice. (A)** Body weight change. **(B)** Epididymal, mesenteric subcutaneous inguinal and retroperitoneal adipose tissue weight (ratio to body weight). NC group, n=10; DSS group, n=10. H&E-stained **(C)** and Oil Red O-stained **(D)** histological sections of liver tissue (200×, scale bar = 0.1 mm) in NC and DSS mice. **(E)** H&E-stained histological sections of epididymal adipose tissue (200×, scale bar = 0.1 mm) in NC and DSS mice. The average size of adipocytes was calculated. NC group, n=10; DSS group, n=10. The level of total triglyceride **(F)** and cholesterol **(G)** in serum in NC and DSS mice. NC group, n=10; DSS group, n=10. The level of total triglyceride **(H)** and cholesterol **(I)** in liver tissue in NC and DSS mice (ratio to total protein). NC group, n=10; DSS group, n=10. NC, the control mice treated with pure drinking water; DSS, the mice were treated with 0.2% (w./v.) DSS added in drinking water. The data are shown as mean ± s.e.m., and Student's t-test (two-tailed) was used to analyze differences of NC vs. DSS.

**Figure S5**

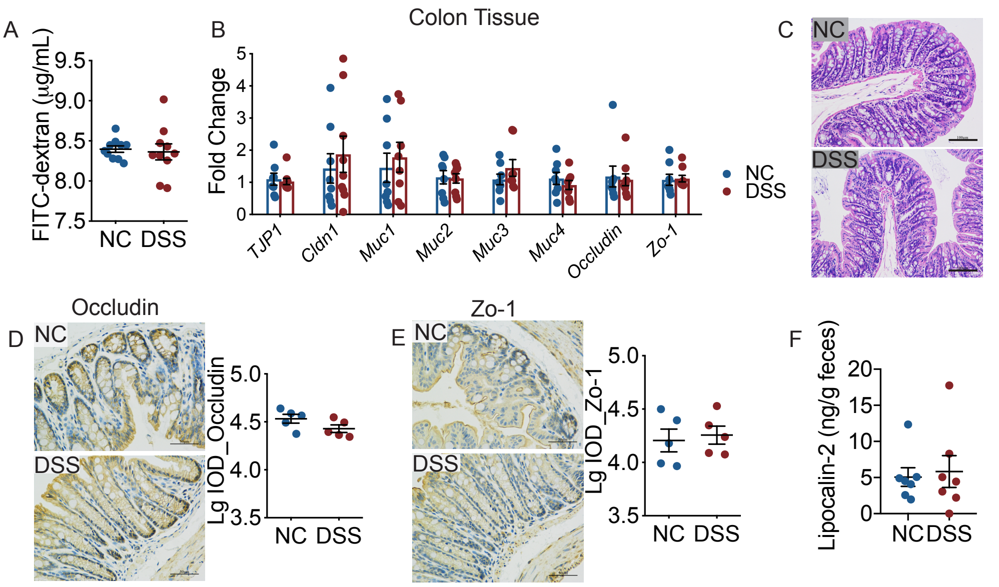


**Fig. S5 0.2% DSS did not impair gut barrier integrity and induce inflammation. (A)** FITC-dextran assessment of the permeability of gut barrier. NC group, n=8; DSS group, n=8. **(B)** Levels of mRNA expression of genes related with gut barrier integrity in the colon tissue, NC group, n=8; DSS group, n=8. **(C)** H&E-stained histological sections of colon tissue (200×, scale bar = 0.1 mm) in NC and DSS mice. Immunohistochemical staining of colon and integrated optical density (IOD) analysis for Occludin **(D)** and Zo-1 **(E)** from NC and DSS mice (n= 5, for each group). **(F)** Fecal lipocalin-2 contents. NC group, n=7; DSS group, n=7. NC, the control mice treated with pure drinking water; DSS, the mice were treated with 0.2% (w./v.) DSS added in drinking water. The data in **(A), (B)** and **(D)-(F)** were expressed as mean ± s.e.m., and Student's t-test (two-tailed) was used to analyze differences between NC vs. DSS. *$P$ <0.05 and **$P$ <0.01.

#### Figure S6

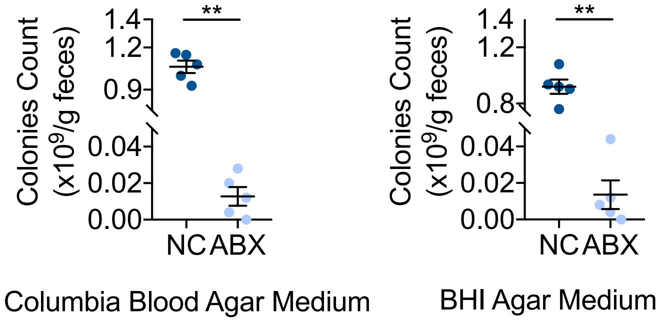


**Fig. S6 The gut microbiota had been depleted by more than 99% in the mice with a cocktail of antibiotics for 5 weeks.** The fecal samples were diluted and spread on the surface of Columbia blood agar medium plates and BHI agar medium plates, and the plates were placed in an anaerobic workstation (80% $N_2$, 10% $CO_2$, and 10% $H_2$ at 37°C) for 48 hours before colonies count was performed. NC group, n=5; ABX group, n=5. NC, the control mice treated with pure drinking water; ABX, the antibiotic-cocktail treated mice with pure water. The data were expressed as mean ± s.e.m., and Student's t-test (two-tailed) was used to analyze differences between NC vs. ABX. $^{**}P < 0.01$.

#### Figure S7

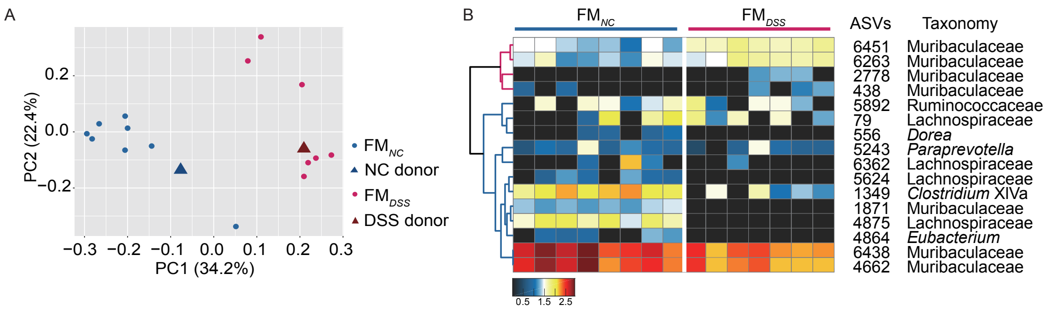


**Fig. S7 The recipient mice developed a gut microbiota more similar to their donor mice. (A)** Principal-coordinate analysis (PCoA) plot of the gut microbial structure of $FM_{NC}$, $FM_{DSS}$, NC donor and DSS donor based on the Bray-Curtis distance. **(B)** Heat map of 16 ASVs in the $FM_{NC}$ and $FM_{DSS}$ mice, which were selected by sPLS-DA model in NC and DSS mice (see Fig. 1H). The cluster tree on the left shows associations between these ASVs as determined by Spearman correlation coefficient based on their relative abundance among all the samples. The heat map on the right shows the relative abundance (log10 transformed) of each ASV in sample of individual mouse. $FM_{NC}$, the antibiotic-cocktail treated mice inoculated with the fecal microbiota from the mice of NC group, and had pure water; $FM_{DSS}$, the antibiotic-cocktail treated mice inoculated with the fecal microbiota from the mice of DSS group and had pure water. $FM_{NC}$ group, n=8; $FM_{DSS}$ group, n=7.

#### Figure S8

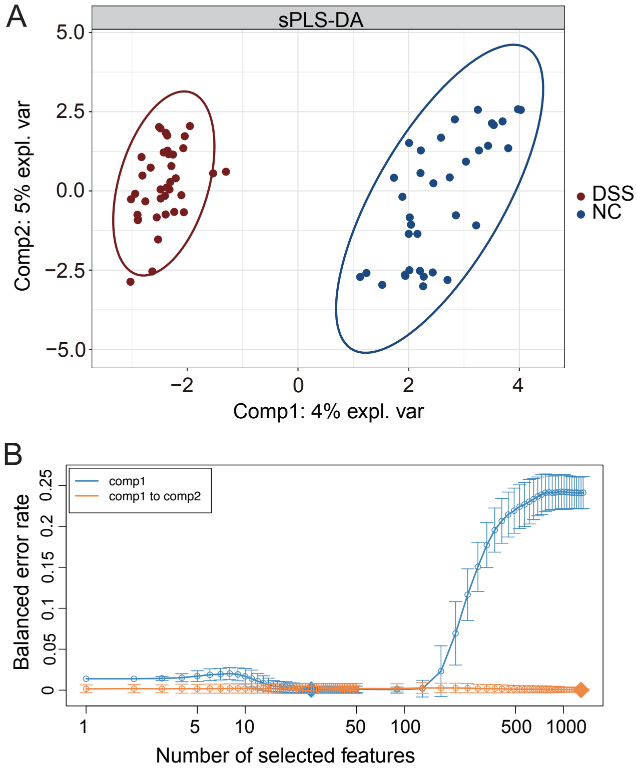


**Fig. S8 Optimal classification performance of the sPLS-DA model of the gut microbial structure in NC and DSS mice. (A)** sPLS-DA plot of the gut microbial structure. **(B)** Error rate of the sPLS-DA model. NC, the control mice treated with pure drinking water; DSS, the mice were treated with 0.2% (w./v.) DSS added in drinking water. The optimal classification performances of the sPLS-DA models were estimated by the perf function using 5-fold cross-validation with the lowest balanced error rate.

#### Figure S9

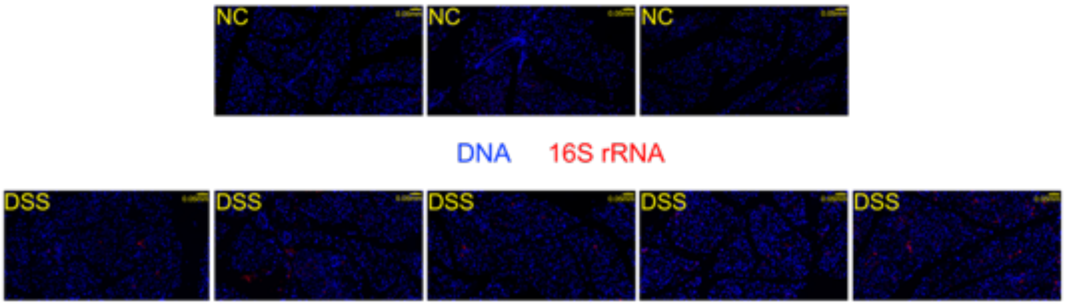


**Fig. S9 Fluorescence in situ hybridization (FISH) of 16S rRNA in the pancreas of antibiotic treated mice.** NC group, n=3; DSS group, n=5. NC, the control mice treated with pure drinking water; DSS, the mice were treated with 0.2% (w./v.) DSS added in drinking water.

#### Figure S10

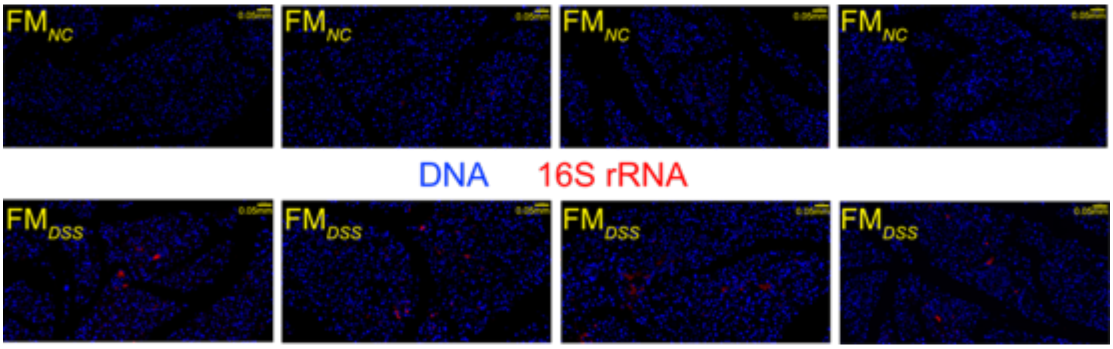


**Fig. S10 Fluorescence in situ hybridization (FISH) of 16S rRNA in the pancreas in fecal microbiota transplanted mice.** $FM_{NC}$ group, n=4; $FM_{DSS}$ group, n=4. $FM_{NC}$, the antibiotic-cocktail treated mice inoculated with the fecal microbiota from the mice of NC group, and had pure water; $FM_{DSS}$, the antibiotic-cocktail treated mice inoculated with the fecal microbiota from the mice of DSS group and had pure water.

#### Figure S11

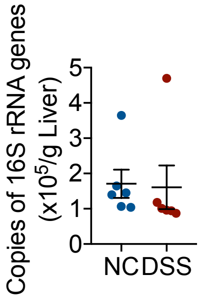


**Fig. S11 The bacterial load in the liver as measured by real-time qPCR of 16S rRNA gene.** NC, n=6; DSS, n=6.

#### Figure S12

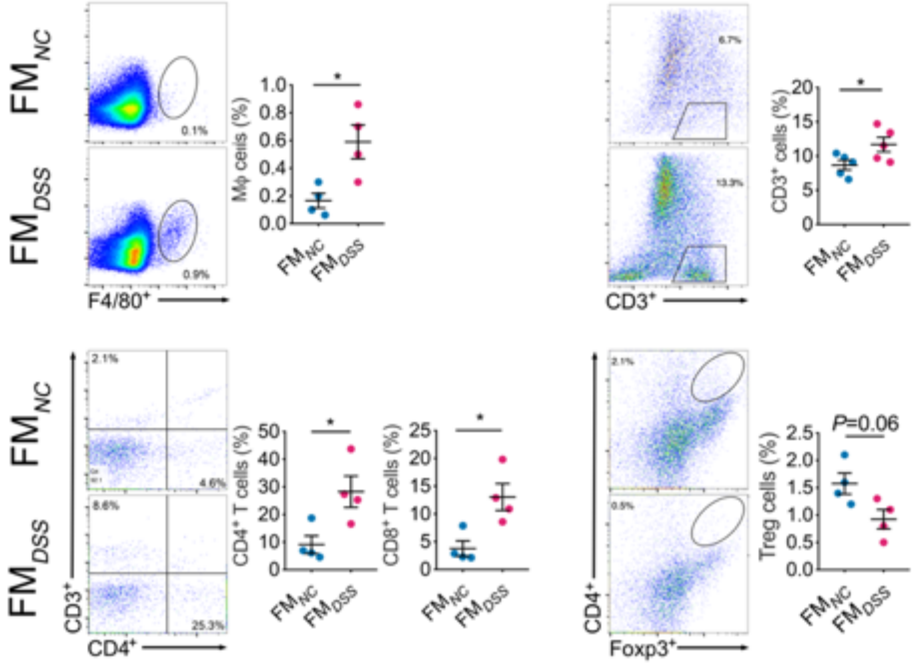


**Fig. S12 Flow cytometry evaluation of subtypes of leukocytes in the pancreas of FM*NC* and FM*DSS* mice.** Data are the frequencies of gated cells (F4/80$^{+}$ macrophage, CD3$^{+}$ cell, CD4$^{+}$ T cell and Foxp3$^{+}$ CD4$^{+}$ cell) among the CD45$^{+}$ cell population per mouse. NC group, n=4-5; DSS group, n=4-5. FM*NC*, the antibiotic-cocktail treated mice inoculated with the fecal microbiota from the mice of NC group, and had pure water; FM*DSS*, the antibiotic-cocktail treated mice inoculated with the fecal microbiota from the mice of DSS group, and had pure water. The data are shown as mean ± s.e.m., and Student's t-test (two-tailed) was used to analyze differences FM*NC* vs. FM*DSS*. *P <0.05 and **P <0.01.

#### Figure S13

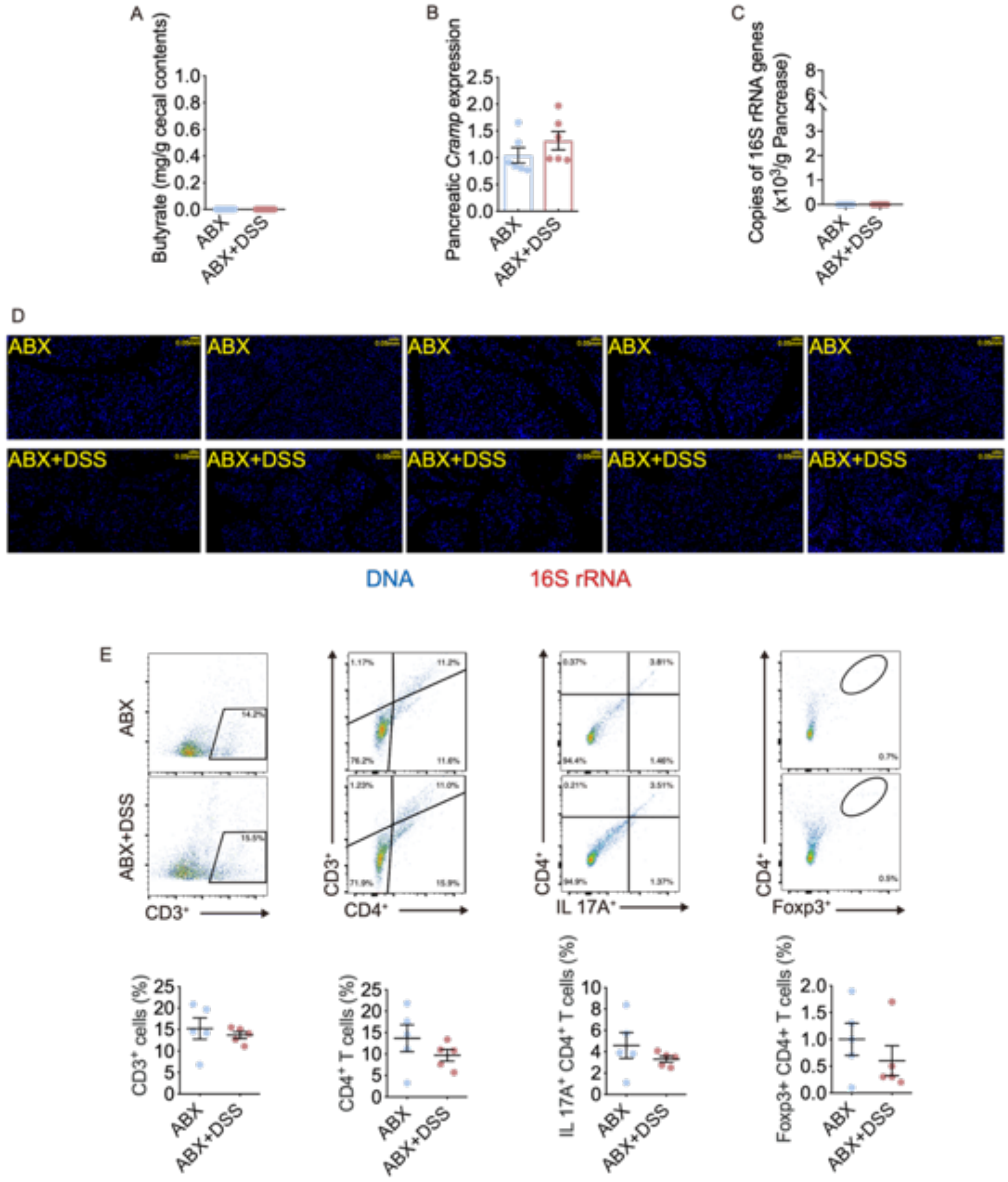


**Fig. S13 0.2% DSS did not enrich bacteria in pacreas and disrupt the immune tolerance in antibiotic treated mice. (A)** The content of butyrate in cecum of mice. NC, n=8; DSS, n=9; ABX, n=8; ABX+DSS, n=8. **(B)** The gene expressions of CRAMP in pancreas of mice. ABX, n=6; ABX+DSS, n=6. The bacterial load in pancreas measured by **(C)** real-time qPCR and **(D)** fluorescence in situ hybridization (FISH) of 16S rRNA. ABX, n=5; ABX+DSS, n=5. **(E)** Flow cytometry evaluation of subtypes of leukocytes in the pancreas of ABX and ABX+DSS mice. Data are the frequency of gated cells ($CD3^+$ cell, $CD4^+$ T cell, IL-$17A^+$ $CD4^+$ cell, and $Foxp3^+$ $CD4^+$ cell) among the $CD45^+$ population of cells per mouse. ABX group, n=5; ABX+DSS group, n=5. The data are shown as mean ± s.e.m., and Student's t-test (two-tailed) was used to analyze differences ABX vs. ABX+DSS.

#### Figure S14

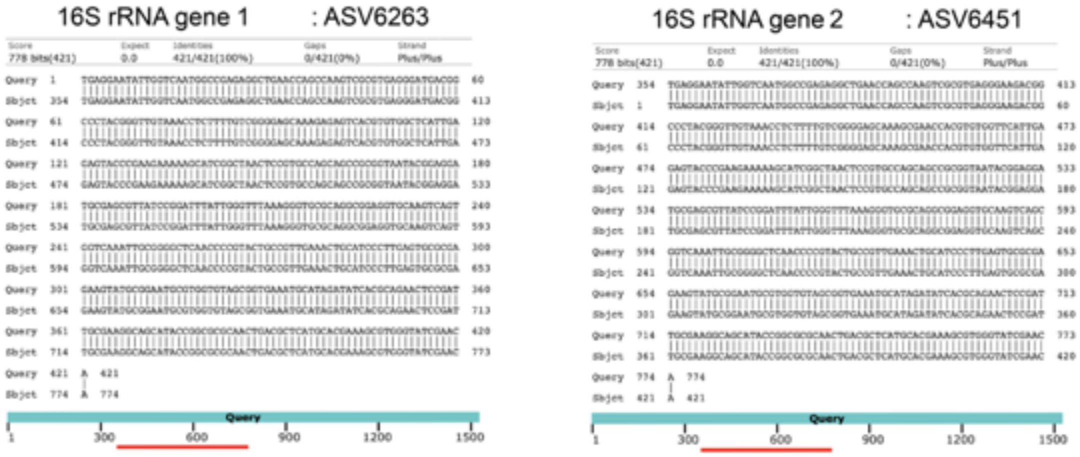


**Fig. S14 Identification of Muribaculaceae strain MF 13079.** Alignment results of 16S rRNA sequence in the genome of MF13079 strain with ASV 6263 and ASV 6451.

#### Figure S15

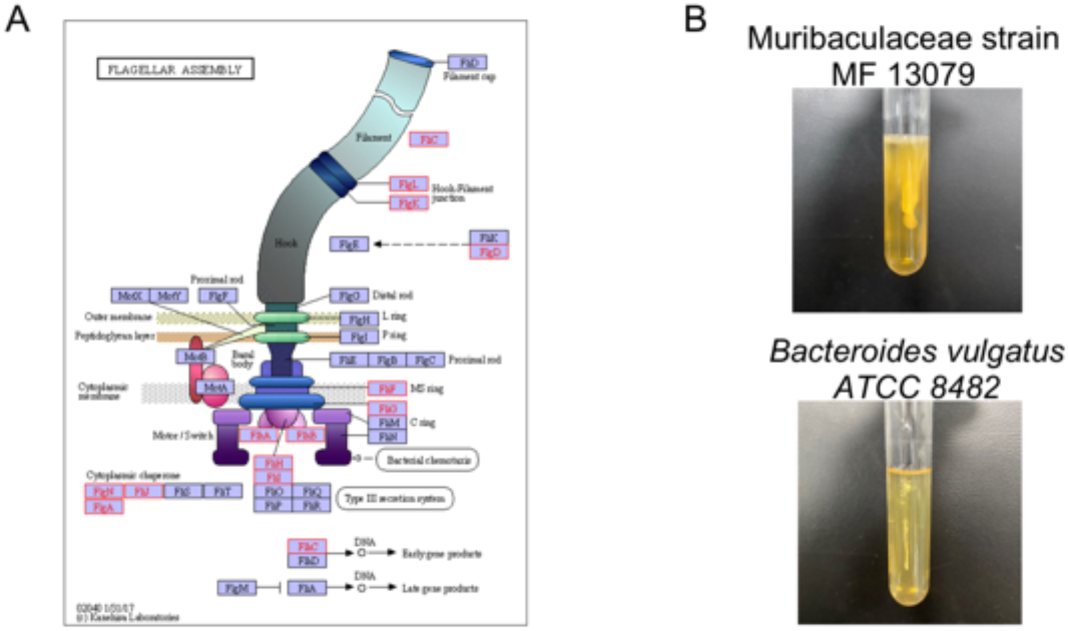


**Fig. S15 Functional annotation of flagella-related genes and motility test of MF13079. (A)** Functional annotation of genes related with flagella-synthesis in the genome of MF13079 using the Kyoto Encyclopedia of Genes and Genomes (KEGG) database. **(B)** Motility test of MF13079 in pre-reduced brain-heart infusion media with 0.3% agar. Motility is designated by growth deviated from the center stab line after 48 h. *Bacteroides vulgatus* ATCC 8482 is non-motile bacterium used as negative control.

**Figure S16**

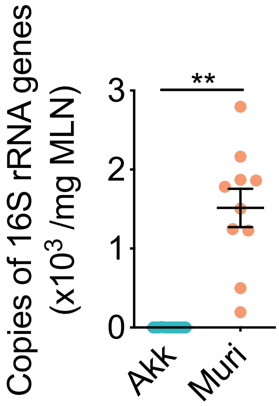


**Fig. S16 Bacteria load in mesenteric lymph nodes (MLN) in Akk and Muri mice.** Akk group, n=12; Muri group, n=10. Muri, the germ-free mice inoculated with the MF 13079 strain, and had pure water; Akk, the germ-free mice inoculated with the Akkermansia muciniphila strain 139 and had pure water. The data were shown as mean ± s.e.m., and Student's t-test (two-tailed) was used to analyze differences between groups. ***P* <0.01.
