## Supplementary Table 1 for "Pathobionts from chemically disrupted gut microbiota induce insulin-dependent diabetes in mice"

### Table S1. Culture medium used for isolation of Muribaculeae

| Medium1 . MPYG medium with 3% FBS | |
|---|---|
| **Component** | **Concentration** |
| Trypticase peptone | 5.00 g/L |
| Peptone | 3.00 g/L |
| Peptone from soya | 2.00 g/L |
| Polypeptone | 1.00 g/L |
| Yeast extract | 10.00 g/L |
| Beef extract | 5.00 g/L |
| Glucose | 5.00 g/L |
| Tween 80 | 0.50 ml/L |
| Maltose | 0.50 g/L |
| Cellobiose | 0.50 g/L |
| Starch, soluble | 0.50 g/L |
| Glycerol | 0.50 ml /L |
| K2HPO4 | 2.00 g/L |
| Cysteine-HCl x H2O | 0.50 g/L |
| Na2S | 0.25 g/L |
| Resazurin | 1.00 mg/L |
| Salt solution (see below) | 40.00 ml/L |
| Trace element(see below) | 10.00 ml/L |
| Vitamin solution(see below) | 10.00 ml/L |
| Haemin solution (see below) | 10.00 ml/L |
| Vitamin K1 solution (see below) | 0.20 ml/L |
| Salt solution(DSMZ Salt solution): | |
| CaCl2 x 2 H2O | 0.25 g/L |
| MgSO4 x 7 H2O | 0.50 g/L |
| K2HPO4 | 1.00 g/L |
| KH2PO4 | 1.00 g/L |
| NaHCO3 | 10.00 g/L |
| NaCl | 2.00 g/L |
| Trace element solution(DSMZ Trace element solution): | |
| Nitrilotriacetic acid | 1.50 g/L |
| MgSO4 x 7 H2O | 3.00 g/L |
| MnSO4 x H2O | 0.50 g/L |
| NaCl | 1.00 g/L |
| FeSO4 x 7 H2O | 0.10 g/L |
| CoSO4 x 7 H2O | 0.18 g/L |
| CaCl2 x 2 H2O | 0.10 g/L |
| ZnSO4 x 7 H2O | 0.18 g/L |
| CuSO4 x 5 H2O | 0.01 g/L |
| KAl(SO4)2 x 12 H2O | 0.02 g/L |
| H3BO3 | 0.01 g/L |
| Na2MoO4 x 2 H2O | 0.01 g/L |
| NiCl2 x 6 H2O | 0.03 g/L |
| Na2SeO3 x 5 H2O | 0.30 mg/L |
| Vitamin solution: | |
| Biotin | 2.00 mg/L |
| Folic acid | 2.00 mg/L |
| Pyridoxine-HCl | 10.00 mg/L |
| Thiamine-HCl x 2 H2O | 5.00 mg/L |
| Riboflavin | 5.00 mg/L |
| Nicotinic acid | 5.00 mg/L |
| D-Ca-pantothenate | 5.00 mg/L |
| Vitamin B12 | 0.10 mg/L |
| p-Aminobenzoic acid | 5.00 mg/L |
| Lipoic acid | 5.00 mg/L |
| Haemin solution | |
| Dissolve 50 mg haemin in 1 ml 1 N NaOH; make up to 100 ml with distilled water. Store refrigerated. | |
| Vitamin K1 solution | |
| Dissolve 0.1 ml of vitamin K1 in 20 ml 95% ethanol and filter sterilize. Store refrigerated in a brown bottle. | |
| Others | |
| Fetal Bovine Serum | 30 mL |

| Medium2 .BHI medium | |
|---|---|
| **Company** | **Product Code** |
| HOPEBIO Co., Ltd | HB8297 |

| Medium3 .BHI medium with mucin | |
|---|---|
| **Company** | **Product Code** |
| HOPEBIO Co., Ltd | HB8297 |
| **Conponents that need to be added** | |
| **Component** | **Concentration** |
| Mucin | 4 g/L |

| Medium4 .BHI medium with DSS | |
|---|---|
| **Company** | **Product Code** |
| HOPEBIO Co., Ltd | HB8297 |
| **Conponents that need to be added** | |
| **Component** | **Concentration** |
| DSS | 1 g/L |

| Medium5 .BHI medium with acetic acid | |
|---|---|
| **Company** | **Product Code** |
| HOPEBIO Co., Ltd | HB8297 |
| **Conponents that need to be added** | |
| **Component** | **Concentration** |
| acetic acid | 33 mM |

| Medium6 .BHI medium with acetic acid, DSS and mucin | |
|---|---|
| **Company** | **Product Code** |
| HOPEBIO Co., Ltd | HB8297 |
| **Conponents that need to be added** | |
| **Component** | **Concentration** |
| acetic acid | 33 mM |
| DSS | 1 g/L |
| Mucin | 4 g/L |

| Medium7 .BHI medium with sulfate | |
|---|---|
| **Company** | **Product Code** |
| HOPEBIO Co., Ltd | HB8297 |
| **Conponents that need to be added** | |
| **Component** | **Concentration** |
| Na2SO4 | 0.1 g/L |
| FeSO4 | 0.05 g/L |
| (NH4)2SO4 | 0.09 g/L |

| Medium8.M2GSC medium (30% CRF) | |
|---|---|
| **Component** | **Concentration** |
| Clarified rumen fluid (CRF) | 300 g/L |
| Casitone | 10 g/L |
| Yeast extract | 2.5 g/L |
| Glucose | 2 g/L |
| Cellobiose | 2 g/L |
| Soluble starch | 2 g/L |
| Cysteine | 1 g/L |
| NaHCO3 | 4 g/L |
| KH2PO4 | 0.45 g/L |
| K2HPO4 | 0.45 g/L |
| (NH4)2SO4 | 0.9 g/L |
| NaCl | 0.9 g/L |
| MgSO4·7H2O | 0.09 g/L |
| CaCl2 | 0.09 g/L |
| Resazurin | 1 g/L |
| Agar | 13 g/L |

| Medium9.M2GSC medium (40% CRF) | |
|---|---|
| **Component** | **Concentration** |
| Clarified rumen fluid (CRF) | 400 g/L |
| Casitone | 10 g/L |
| Yeast extract | 2.5 g/L |
| Glucose | 2 g/L |
| Cellobiose | 2 g/L |
| Soluble starch | 2 g/L |
| Cysteine | 1 g/L |
| NaHCO3 | 4 g/L |
| KH2PO4 | 0.45 g/L |
| K2HPO4 | 0.45 g/L |
| (NH4)2SO4 | 0.9 g/L |
| NaCl | 0.9 g/L |
| MgSO4·7H2O | 0.09 g/L |
| CaCl2 | 0.09 g/L |
| Resazurin | 1 g/L |
| Agar | 13 g/L |

| Medium10.M2GSC medium with mucin | |
|---|---|
| **Component** | **Concentration** |
| Clarified rumen fluid (CRF) | 300 g/L |
| Casitone | 10 g/L |
| Yeast extract | 2.5 g/L |
| Glucose | 2 g/L |
| Cellobiose | 2 g/L |
| Soluble starch | 2 g/L |
| Cysteine | 1 g/L |
| NaHCO3 | 4 g/L |
| KH2PO4 | 0.45 g/L |
| K2HPO4 | 0.45 g/L |
| (NH4)2SO4 | 0.9 g/L |
| NaCl | 0.9 g/L |
| MgSO4·7H2O | 0.09 g/L |
| CaCl2 | 0.09 g/L |
| Resazurin | 1 g/L |
| Agar | 13 g/L |
| **Conponents that need to be added** | |
| **Component** | **Concentration** |
| Mucin | 4 g/L |

| Medium11.M2GSC medium with DSS | |
|---|---|
| **Component** | **Concentration** |
| Clarified rumen fluid (CRF) | 300 g/L |
| Casitone | 10 g/L |
| Yeast extract | 2.5 g/L |
| Glucose | 2 g/L |
| Cellobiose | 2 g/L |
| Soluble starch | 2 g/L |
| Cysteine | 1 g/L |
| NaHCO3 | 4 g/L |
| KH2PO4 | 0.45 g/L |
| K2HPO4 | 0.45 g/L |
| (NH4)2SO4 | 0.9 g/L |
| NaCl | 0.9 g/L |
| MgSO4·7H2O | 0.09 g/L |
| CaCl2 | 0.09 g/L |
| Resazurin | 1 g/L |
| Agar | 13 g/L |
| **Conponents that need to be added** | |
| **Component** | **Concentration** |
| DSS | 1 g/L |

| Medium12.M2GSC medium with acedic acid | |
|---|---|
| **Component** | **Concentration** |
| Clarified rumen fluid (CRF) | 400 g/L |
| Casitone | 10 g/L |
| Yeast extract | 2.5 g/L |
| Glucose | 2 g/L |
| Cellobiose | 2 g/L |
| Soluble starch | 2 g/L |
| Cysteine | 1 g/L |
| NaHCO3 | 4 g/L |
| KH2PO4 | 0.45 g/L |
| K2HPO4 | 0.45 g/L |
| (NH4)2SO4 | 0.9 g/L |
| NaCl | 0.9 g/L |
| MgSO4·7H2O | 0.09 g/L |
| CaCl2 | 0.09 g/L |
| Resazurin | 1 g/L |
| Agar | 13 g/L |
| **Conponents that need to be added** | |
| **Component** | **Concentration** |
| acetic acid | 33 mM |

| Medium13.M2GSC medium with acetic acid, DSS and mucin | |
|---|---|
| **Component** | **Concentration** |
| Clarified rumen fluid (CRF) | 300 g/L |
| Casitone | 10 g/L |
| Yeast extract | 2.5 g/L |
| Glucose | 2 g/L |
| Cellobiose | 2 g/L |
| Soluble starch | 2 g/L |
| Cysteine | 1 g/L |
| NaHCO3 | 4 g/L |
| KH2PO4 | 0.45 g/L |
| K2HPO4 | 0.45 g/L |
| (NH4)2SO4 | 0.9 g/L |
| NaCl | 0.9 g/L |
| MgSO4·7H2O | 0.09 g/L |
| CaCl2 | 0.09 g/L |
| Resazurin | 1 g/L |
| Agar | 13 g/L |
| **Conponents that need to be added** | |
| **Component** | **Concentration** |
| acetic acid | 33 mM |
| DSS | 1 g/L |
| Mucin | 4 g/L |

| Medium14.M2GSC medium with sulfate | |
|---|---|
| **Component** | **Concentration** |
| Clarified rumen fluid (CRF) | 400 g/L |
| Casitone | 10 g/L |
| Yeast extract | 2.5 g/L |
| Glucose | 2 g/L |
| Cellobiose | 2 g/L |
| Soluble starch | 2 g/L |
| Cysteine | 1 g/L |
| NaHCO3 | 4 g/L |
| KH2PO4 | 0.45 g/L |
| K2HPO4 | 0.45 g/L |
| (NH4)2SO4 | 0.9 g/L |
| NaCl | 0.9 g/L |
| MgSO4·7H2O | 0.09 g/L |
| CaCl2 | 0.09 g/L |
| Resazurin | 1 g/L |
| Agar | 13 g/L |
| **Conponents that need to be added** | |
| **Component** | **Concentration** |
| Na2SO4 | 0.1 g/L |
| FeSO4 | 0.05 g/L |
| (NH4)2SO4 | 0.09 g/L |

| Medium15.mGAM medium | |
|---|---|
| **Company** | **Product Code** |
| HOPEBIO Co., Ltd | HB8462 |

| Medium16.mGAM medium with mucin | |
|---|---|
| **Company** | **Product Code** |
| HOPEBIO Co., Ltd | HB8462 |
| **Conponents that need to be added** | |
| **Component** | **Concentration** |
| Mucin | 4 g/L |

| Medium17.mGAM medium with DSS | |
|---|---|
| **Company** | **Product Code** |
| HOPEBIO Co., Ltd | HB8462 |
| **Conponents that need to be added** | |
| **Component** | **Concentration** |
| DSS | 1 g/L |

| Medium18.mGAM medium with acetic acid | |
|---|---|
| **Company** | **Product Code** |
| HOPEBIO Co., Ltd | HB8462 |
| **Conponents that need to be added** | |
| **Component** | **Concentration** |
| acetic acid | 33 mM |

| Medium19.mGAM medium with acetic acid, DSS and mucin | |
|---|---|
| **Company** | **Product Code** |
| HOPEBIO Co., Ltd | HB8462 |
| **Conponents that need to be added** | |
| **Component** | **Concentration** |
| acetic acid | 33 mM |
| DSS | 1 g/L |
| Mucin | 4 g/L |

| Medium20.mGAM medium with sulfate | |
|---|---|
| **Company** | **Product Code** |
| HOPEBIO Co., Ltd | HB8462 |
| **Conponents that need to be added** | |
| **Component** | **Concentration** |
| Na2SO4 | 0.1 g/L |
| FeSO4 | 0.05 g/L |
| (NH4)2SO4 | 0.09 g/L |

| Medium21.mGAM medium with SCFAs | |
|---|---|
| **Company** | **Product Code** |
| HOPEBIO Co., Ltd | HB8462 |
| **Conponents that need to be added** | |
| **Component** | **Concentration** |
| acetic acid | 33 mM |
| propionic acid | 8 mM |
| butyric acid | 4 mM |

| Medium22.A2 medium | |
|---|---|
| **Component** | **Concentration** |
| BHI medium(HB8297, HOPEBIO Co., Ltd) | 18.5 g/L |
| Yeast extract | 5 g/L |
| TSB | 15 g/L |
| K2HPO4 | 2.5 g/L |
| hemin | 1 mg/L |
| glucose | 1 g/L |
| L-cysteine | 0.5 g/L |
| Na2CO3 | 0.4 g/L |
| VK3 | 1 mg/L |
| Fetal Bovine Serum | 30 mL/L |

| Medium23. A2 medium with mucin | |
|---|---|
| **Component** | **Concentration** |
| BHI medium(HB8297, HOPEBIO Co., Ltd) | 18.5 g/L |
| Yeast extract | 5 g/L |
| TSB | 15 g/L |
| K2HPO4 | 2.5 g/L |
| hemin | 1 mg/L |
| glucose | 1 g/L |
| L-cysteine | 0.5 g/L |
| Na2CO3 | 0.4 g/L |
| VK3 | 1 mg/L |
| Fetal Bovine Serum | 30 mL/L |
| **Conponents that need to be added** | |
| **Component** | **Concentration** |
| Mucin | 4 g/L |

| Medium24. A2 medium with acetic acid | |
|---|---|
| **Component** | **Concentration** |
| BHI medium(HB8297, HOPEBIO Co., Ltd) | 18.5 g/L |
| Yeast extract | 5 g/L |
| TSB | 15 g/L |
| K2HPO4 | 2.5 g/L |
| hemin | 1 mg/L |
| glucose | 1 g/L |
| L-cysteine | 0.5 g/L |
| Na2CO3 | 0.4 g/L |
| VK3 | 1 mg/L |
| Fetal Bovine Serum | 30 mL/L |
| **Conponents that need to be added** | |
| **Component** | **Concentration** |
| acetic acid | 33 mM |

| Medium25.A2 medium with DSS | |
|---|---|
| **Component** | **Concentration** |
| BHI medium(HB8297, HOPEBIO Co., Ltd) | 18.5 g/L |
| Yeast extract | 5 g/L |
| TSB | 15 g/L |
| K2HPO4 | 2.5 g/L |
| hemin | 1 mg/L |
| glucose | 1 g/L |
| L-cysteine | 0.5 g/L |
| Na2CO3 | 0.4 g/L |
| VK3 | 1 mg/L |
| Fetal Bovine Serum | 30 mL/L |
| **Conponents that need to be added** | |
| **Component** | **Concentration** |
| DSS | 1 g/L |

| Medium26. A2 medium with acetic acid, DSS and mucin | |
|---|---|
| **Component** | **Concentration** |
| BHI medium(HB8297, HOPEBIO Co., Ltd) | 18.5 g/L |
| Yeast extract | 5 g/L |
| TSB | 15 g/L |
| K2HPO4 | 2.5 g/L |
| hemin | 1 mg/L |
| glucose | 1 g/L |
| L-cysteine | 0.5 g/L |
| Na2CO3 | 0.4 g/L |
| VK3 | 1 mg/L |
| Fetal Bovine Serum | 30 mL/L |
| **Conponents that need to be added** | |
| **Component** | **Concentration** |
| acetic acid | 33 mM |
| DSS | 1 g/L |
| Mucin | 4 g/L |

| Medium27. A2 medium with sulfate | |
|---|---|
| **Component** | **Concentration** |
| BHI medium(HB8297, HOPEBIO Co., Ltd) | 18.5 g/L |
| Yeast extract | 5 g/L |
| TSB | 15 g/L |
| K2HPO4 | 2.5 g/L |
| hemin | 1 mg/L |
| glucose | 1 g/L |
| L-cysteine | 0.5 g/L |
| Na2CO3 | 0.4 g/L |
| VK3 | 1 mg/L |
| Fetal Bovine Serum | 30 mL/L |
| **Conponents that need to be added** | |
| **Component** | **Concentration** |
| Na2SO4 | 0.1 g/L |
| FeSO4 | 0.05 g/L |
| (NH4)2SO4 | 0.09 g/L |

| Medium28. A2 medium with SCFAs | |
|---|---|
| **Component** | **Concentration** |
| BHI medium(HB8297, HOPEBIO Co., Ltd) | 18.5 g/L |
| Yeast extract | 5 g/L |
| TSB | 15 g/L |
| K2HPO4 | 2.5 g/L |
| hemin | 1 mg/L |
| glucose | 1 g/L |
| L-cysteine | 0.5 g/L |
| Na2CO3 | 0.4 g/L |
| VK3 | 1 mg/L |
| Fetal Bovine Serum | 30 mL/L |
| **Conponents that need to be added** | |
| **Component** | **Concentration** |
| acetic acid | 33 mM |
| propionic acid | 8 mM |
| butyric acid | 4 mM |

| Medium29. BHI medium with SCFAs | |
|---|---|
| **Company** | **Product Code** |
| HOPEBIO Co., Ltd | HB8297 |
| **Conponents that need to be added** | |
| **Component** | **Concentration** |
| acetic acid | 33 mM |
| propionic acid | 8 mM |
| butyric acid | 4 mM |

| Medium30.Columbia Blood Agar medium | |
|---|---|
| **Company** | **Product Code** |
| HOPEBIO Co., Ltd | HB0124 |
| **Conponents that need to be added** | |
| **Component** | **Concentration** |
| Off fiber sheep blood | 50 mL/L |

| Medium31. Columbia Blood Agar medium with DSS | |
|---|---|
| **Company** | **Product Code** |
| HOPEBIO Co., Ltd | HB0124 |
| **Conponents that need to be added** | |
| **Component** | **Concentration** |
| Off fiber sheep blood | 50 mL/L |
| DSS | 1 g/L |

| Medium32. Columbia Blood Agar medium with mucin | |
|---|---|
| **Company** | **Product Code** |
| HOPEBIO Co., Ltd | HB0124 |
| **Conponents that need to be added** | |
| **Component** | **Concentration** |
| Off fiber sheep blood | 50 mL/L |
| mucin | 4 g/L |

| Medium33. Columbia Blood Agar medium with mucin and DSS | |
|---|---|
| **Company** | **Product Code** |
| HOPEBIO Co., Ltd | HB0124 |
| **Conponents that need to be added** | |
| **Component** | **Concentration** |
| Off fiber sheep blood | 50 mL/L |
| mucin | 4 g/L |
| DSS | 1 g/L |

| Medium34. MPYG medium | |
|---|---|
| **Component** | **Concentration** |
| Trypticase peptone | 5.00 g/L |
| Peptone | 3.00 g/L |
| Peptone from soya | 2.00 g/L |
| Polypeptone | 1.00 g/L |
| Yeast extract | 10.00 g/L |
| Beef extract | 5.00 g/L |
| Glucose | 5.00 g/L |
| Tween 80 | 0.50 ml/L |
| Maltose | 0.50 g/L |
| Cellobiose | 0.50 g/L |
| Starch, soluble | 0.50 g/L |
| Glycerol | 0.50 ml /L |
| K2HPO4 | 2.00 g/L |
| Cysteine-HCl x H2O | 0.50 g/L |
| Na2S | 0.25 g/L |
| Resazurin | 1.00 mg/L |
| Salt solution (see below) | 40.00 ml/L |
| Trace element(see below) | 10.00 ml/L |
| Vitamin solution(see below) | 10.00 ml/L |
| Haemin solution (see below) | 10.00 ml/L |
| Vitamin K1 solution (see below) | 0.20 ml/L |
| Salt solution(DSMZ Salt solution): | |
| CaCl2 x 2 H2O | 0.25 g/L |
| MgSO4 x 7 H2O | 0.50 g/L |
| K2HPO4 | 1.00 g/L |
| KH2PO4 | 1.00 g/L |
| NaHCO3 | 10.00 g/L |
| NaCl | 2.00 g/L |
| Trace element solution(DSMZ Trace element solution): | |
| Nitrilotriacetic acid | 1.50 g/L |
| MgSO4 x 7 H2O | 3.00 g/L |
| MnSO4 x H2O | 0.50 g/L |
| NaCl | 1.00 g/L |
| FeSO4 x 7 H2O | 0.10 g/L |
| CoSO4 x 7 H2O | 0.18 g/L |
| CaCl2 x 2 H2O | 0.10 g/L |
| ZnSO4 x 7 H2O | 0.18 g/L |
| CuSO4 x 5 H2O | 0.01 g/L |
| KAl(SO4)2 x 12 H2O | 0.02 g/L |
| H3BO3 | 0.01 g/L |
| Na2MoO4 x 2 H2O | 0.01 g/L |
| NiCl2 x 6 H2O | 0.03 g/L |
| Na2SeO3 x 5 H2O | 0.30 mg/L |
| Vitamin solution: | |
| Biotin | 2.00 mg/L |
| Folic acid | 2.00 mg/L |
| Pyridoxine-HCl | 10.00 mg/L |
| Thiamine-HCl x 2 H2O | 5.00 mg/L |
| Riboflavin | 5.00 mg/L |
| Nicotinic acid | 5.00 mg/L |
| D-Ca-pantothenate | 5.00 mg/L |
| Vitamin B12 | 0.10 mg/L |
| p-Aminobenzoic acid | 5.00 mg/L |
| Lipoic acid | 5.00 mg/L |
| Haemin solution | |
| Dissolve 50 mg haemin in 1 ml 1 N NaOH; make up to 100 ml with distilled water. Store refrigerated. | |
| Vitamin K1 solution | |
| Dissolve 0.1 ml of vitamin K1 in 20 ml 95% ethanol and filter sterilize. Store refrigerated in a brown bottle. | |

| Medium35. MPYG medium with mucin | |
|---|---|
| **Component** | **Concentration** |
| Trypticase peptone | 5.00 g/L |
| Peptone | 3.00 g/L |
| Peptone from soya | 2.00 g/L |
| Polypeptone | 1.00 g/L |
| Yeast extract | 10.00 g/L |
| Beef extract | 5.00 g/L |
| Glucose | 5.00 g/L |
| Tween 80 | 0.50 ml/L |
| Maltose | 0.50 g/L |
| Cellobiose | 0.50 g/L |
| Starch, soluble | 0.50 g/L |
| Glycerol | 0.50 ml /L |
| K2HPO4 | 2.00 g/L |
| Cysteine-HCl x H2O | 0.50 g/L |
| Na2S | 0.25 g/L |
| Resazurin | 1.00 mg/L |
| Salt solution (see below) | 40.00 ml/L |
| Trace element(see below) | 10.00 ml/L |
| Vitamin solution(see below) | 10.00 ml/L |
| Haemin solution (see below) | 10.00 ml/L |
| Vitamin K1 solution (see below) | 0.20 ml/L |
| Salt solution(DSMZ Salt solution): | |
| CaCl2 x 2 H2O | 0.25 g/L |
| MgSO4 x 7 H2O | 0.50 g/L |
| K2HPO4 | 1.00 g/L |
| KH2PO4 | 1.00 g/L |
| NaHCO3 | 10.00 g/L |
| NaCl | 2.00 g/L |
| Trace element solution(DSMZ Trace element solution): | |
| Nitrilotriacetic acid | 1.50 g/L |
| MgSO4 x 7 H2O | 3.00 g/L |
| MnSO4 x H2O | 0.50 g/L |
| NaCl | 1.00 g/L |
| FeSO4 x 7 H2O | 0.10 g/L |
| CoSO4 x 7 H2O | 0.18 g/L |
| CaCl2 x 2 H2O | 0.10 g/L |
| ZnSO4 x 7 H2O | 0.18 g/L |
| CuSO4 x 5 H2O | 0.01 g/L |
| KAl(SO4)2 x 12 H2O | 0.02 g/L |
| H3BO3 | 0.01 g/L |
| Na2MoO4 x 2 H2O | 0.01 g/L |
| NiCl2 x 6 H2O | 0.03 g/L |
| Na2SeO3 x 5 H2O | 0.30 mg/L |
| Vitamin solution: | |
| Biotin | 2.00 mg/L |
| Folic acid | 2.00 mg/L |
| Pyridoxine-HCl | 10.00 mg/L |
| Thiamine-HCl x 2 H2O | 5.00 mg/L |
| Riboflavin | 5.00 mg/L |
| Nicotinic acid | 5.00 mg/L |
| D-Ca-pantothenate | 5.00 mg/L |
| Vitamin B12 | 0.10 mg/L |
| p-Aminobenzoic acid | 5.00 mg/L |
| Lipoic acid | 5.00 mg/L |
| Haemin solution | |
| Dissolve 50 mg haemin in 1 ml 1 N NaOH; make up to 100 ml with distilled water. Store refrigerated. | |
| Vitamin K1 solution | |
| Dissolve 0.1 ml of vitamin K1 in 20 ml 95% ethanol and filter sterilize. Store refrigerated in a brown bottle. | |
| Others | |
| mucin | 4 g/L |

| Medium36. MPYG medium with DSS | |
|---|---|
| **ComponentAAB3:AC61** | **Concentration** |
| Trypticase peptone | 5.00 g/L |
| Peptone | 3.00 g/L |
| Peptone from soya | 2.00 g/L |
| Polypeptone | 1.00 g/L |
| Yeast extract | 10.00 g/L |
| Beef extract | 5.00 g/L |
| Glucose | 5.00 g/L |
| Tween 80 | 0.50 ml/L |
| Maltose | 0.50 g/L |
| Cellobiose | 0.50 g/L |
| Starch, soluble | 0.50 g/L |
| Glycerol | 0.50 ml /L |
| K2HPO4 | 2.00 g/L |
| Cysteine-HCl x H2O | 0.50 g/L |
| Na2S | 0.25 g/L |
| Resazurin | 1.00 mg/L |
| Salt solution (see below) | 40.00 ml/L |
| Trace element(see below) | 10.00 ml/L |
| Vitamin solution(see below) | 10.00 ml/L |
| Haemin solution (see below) | 10.00 ml/L |
| Vitamin K1 solution (see below) | 0.20 ml/L |
| Salt solution(DSMZ Salt solution): | |
| CaCl2 x 2 H2O | 0.25 g/L |
| MgSO4 x 7 H2O | 0.50 g/L |
| K2HPO4 | 1.00 g/L |
| KH2PO4 | 1.00 g/L |
| NaHCO3 | 10.00 g/L |
| NaCl | 2.00 g/L |
| Trace element solution(DSMZ Trace element solution): | |
| Nitrilotriacetic acid | 1.50 g/L |
| MgSO4 x 7 H2O | 3.00 g/L |
| MnSO4 x H2O | 0.50 g/L |
| NaCl | 1.00 g/L |
| FeSO4 x 7 H2O | 0.10 g/L |
| CoSO4 x 7 H2O | 0.18 g/L |
| CaCl2 x 2 H2O | 0.10 g/L |
| ZnSO4 x 7 H2O | 0.18 g/L |
| CuSO4 x 5 H2O | 0.01 g/L |
| KAl(SO4)2 x 12 H2O | 0.02 g/L |
| H3BO3 | 0.01 g/L |
| Na2MoO4 x 2 H2O | 0.01 g/L |
| NiCl2 x 6 H2O | 0.03 g/L |
| Na2SeO3 x 5 H2O | 0.30 mg/L |
| Vitamin solution: | |
| Biotin | 2.00 mg/L |
| Folic acid | 2.00 mg/L |
| Pyridoxine-HCl | 10.00 mg/L |
| Thiamine-HCl x 2 H2O | 5.00 mg/L |
| Riboflavin | 5.00 mg/L |
| Nicotinic acid | 5.00 mg/L |
| D-Ca-pantothenate | 5.00 mg/L |
| Vitamin B12 | 0.10 mg/L |
| p-Aminobenzoic acid | 5.00 mg/L |
| Lipoic acid | 5.00 mg/L |
| Haemin solution | |
| Dissolve 50 mg haemin in 1 ml 1 N NaOH; make up to 100 ml with distilled water. Store refrigerated. | |
| Vitamin K1 solution | |
| Dissolve 0.1 ml of vitamin K1 in 20 ml 95% ethanol and filter sterilize. Store refrigerated in a brown bottle. | |
| Others | |
| DSS | 1 g/L |

| Medium37. MPYG medium with acetic acid | |
|---|---|
| **Component** | **Concentration** |
| Trypticase peptone | 5.00 g/L |
| Peptone | 3.00 g/L |
| Peptone from soya | 2.00 g/L |
| Polypeptone | 1.00 g/L |
| Yeast extract | 10.00 g/L |
| Beef extract | 5.00 g/L |
| Glucose | 5.00 g/L |
| Tween 80 | 0.50 ml/L |
| Maltose | 0.50 g/L |
| Cellobiose | 0.50 g/L |
| Starch, soluble | 0.50 g/L |
| Glycerol | 0.50 ml /L |
| K2HPO4 | 2.00 g/L |
| Cysteine-HCl x H2O | 0.50 g/L |
| Na2S | 0.25 g/L |
| Resazurin | 1.00 mg/L |
| Salt solution (see below) | 40.00 ml/L |
| Trace element(see below) | 10.00 ml/L |
| Vitamin solution(see below) | 10.00 ml/L |
| Haemin solution (see below) | 10.00 ml/L |
| Vitamin K1 solution (see below) | 0.20 ml/L |
| Salt solution(DSMZ Salt solution): | |
| CaCl2 x 2 H2O | 0.25 g/L |
| MgSO4 x 7 H2O | 0.50 g/L |
| K2HPO4 | 1.00 g/L |
| KH2PO4 | 1.00 g/L |
| NaHCO3 | 10.00 g/L |
| NaCl | 2.00 g/L |
| Trace element solution(DSMZ Trace element solution): | |
| Nitrilotriacetic acid | 1.50 g/L |
| MgSO4 x 7 H2O | 3.00 g/L |
| MnSO4 x H2O | 0.50 g/L |
| NaCl | 1.00 g/L |
| FeSO4 x 7 H2O | 0.10 g/L |
| CoSO4 x 7 H2O | 0.18 g/L |
| CaCl2 x 2 H2O | 0.10 g/L |
| ZnSO4 x 7 H2O | 0.18 g/L |
| CuSO4 x 5 H2O | 0.01 g/L |
| KAl(SO4)2 x 12 H2O | 0.02 g/L |
| H3BO3 | 0.01 g/L |
| Na2MoO4 x 2 H2O | 0.01 g/L |
| NiCl2 x 6 H2O | 0.03 g/L |
| Na2SeO3 x 5 H2O | 0.30 mg/L |
| Vitamin solution: | |
| Biotin | 2.00 mg/L |
| Folic acid | 2.00 mg/L |
| Pyridoxine-HCl | 10.00 mg/L |
| Thiamine-HCl x 2 H2O | 5.00 mg/L |
| Riboflavin | 5.00 mg/L |
| Nicotinic acid | 5.00 mg/L |
| D-Ca-pantothenate | 5.00 mg/L |
| Vitamin B12 | 0.10 mg/L |
| p-Aminobenzoic acid | 5.00 mg/L |
| Lipoic acid | 5.00 mg/L |
| Haemin solution | |
| Dissolve 50 mg haemin in 1 ml 1 N NaOH; make up to 100 ml with distilled water. Store refrigerated. | |
| Vitamin K1 solution | |
| Dissolve 0.1 ml of vitamin K1 in 20 ml 95% ethanol and filter sterilize. Store refrigerated in a brown bottle. | |
| Others | |
| acetic acid | 33 mM |

| Medium38. MPYG medium with mucin, DSS and acetic acid | |
|---|---|
| **Component** | **Concentration** |
| Trypticase peptone | 5.00 g/L |
| Peptone | 3.00 g/L |
| Peptone from soya | 2.00 g/L |
| Polypeptone | 1.00 g/L |
| Yeast extract | 10.00 g/L |
| Beef extract | 5.00 g/L |
| Glucose | 5.00 g/L |
| Tween 80 | 0.50 ml/L |
| Maltose | 0.50 g/L |
| Cellobiose | 0.50 g/L |
| Starch, soluble | 0.50 g/L |
| Glycerol | 0.50 ml /L |
| K2HPO4 | 2.00 g/L |
| Cysteine-HCl x H2O | 0.50 g/L |
| Na2S | 0.25 g/L |
| Resazurin | 1.00 mg/L |
| Salt solution (see below) | 40.00 ml/L |
| Trace element(see below) | 10.00 ml/L |
| Vitamin solution(see below) | 10.00 ml/L |
| Haemin solution (see below) | 10.00 ml/L |
| Vitamin K1 solution (see below) | 0.20 ml/L |
| Salt solution(DSMZ Salt solution): | |
| CaCl2 x 2 H2O | 0.25 g/L |
| MgSO4 x 7 H2O | 0.50 g/L |
| K2HPO4 | 1.00 g/L |
| KH2PO4 | 1.00 g/L |
| NaHCO3 | 10.00 g/L |
| NaCl | 2.00 g/L |
| Trace element solution(DSMZ Trace element solution): | |
| Nitrilotriacetic acid | 1.50 g/L |
| MgSO4 x 7 H2O | 3.00 g/L |
| MnSO4 x H2O | 0.50 g/L |
| NaCl | 1.00 g/L |
| FeSO4 x 7 H2O | 0.10 g/L |
| CoSO4 x 7 H2O | 0.18 g/L |
| CaCl2 x 2 H2O | 0.10 g/L |
| ZnSO4 x 7 H2O | 0.18 g/L |
| CuSO4 x 5 H2O | 0.01 g/L |
| KAl(SO4)2 x 12 H2O | 0.02 g/L |
| H3BO3 | 0.01 g/L |
| Na2MoO4 x 2 H2O | 0.01 g/L |
| NiCl2 x 6 H2O | 0.03 g/L |
| Na2SeO3 x 5 H2O | 0.30 mg/L |
| Vitamin solution: | |
| Biotin | 2.00 mg/L |
| Folic acid | 2.00 mg/L |
| Pyridoxine-HCl | 10.00 mg/L |
| Thiamine-HCl x 2 H2O | 5.00 mg/L |
| Riboflavin | 5.00 mg/L |
| Nicotinic acid | 5.00 mg/L |
| D-Ca-pantothenate | 5.00 mg/L |
| Vitamin B12 | 0.10 mg/L |
| p-Aminobenzoic acid | 5.00 mg/L |
| Lipoic acid | 5.00 mg/L |
| Haemin solution | |
| Dissolve 50 mg haemin in 1 ml 1 N NaOH; make up to 100 ml with distilled water. Store refrigerated. | |
| Vitamin K1 solution | |
| Dissolve 0.1 ml of vitamin K1 in 20 ml 95% ethanol and filter sterilize. Store refrigerated in a brown bottle. | |
| Others | |
| mucin | 4 g/L |
| DSS | 1 g/L |
| acetic acid | 33 mM |

| Medium39. MPYG medium with sulfate | |
|---|---|
| **Component** | **Concentration** |
| Trypticase peptone | 5.00 g/L |
| Peptone | 3.00 g/L |
| Peptone from soya | 2.00 g/L |
| Polypeptone | 1.00 g/L |
| Yeast extract | 10.00 g/L |
| Beef extract | 5.00 g/L |
| Glucose | 5.00 g/L |
| Tween 80 | 0.50 ml/L |
| Maltose | 0.50 g/L |
| Cellobiose | 0.50 g/L |
| Starch, soluble | 0.50 g/L |
| Glycerol | 0.50 ml /L |
| K2HPO4 | 2.00 g/L |
| Cysteine-HCl x H2O | 0.50 g/L |
| Na2S | 0.25 g/L |
| Resazurin | 1.00 mg/L |
| Salt solution (see below) | 40.00 ml/L |
| Trace element(see below) | 10.00 ml/L |
| Vitamin solution(see below) | 10.00 ml/L |
| Haemin solution (see below) | 10.00 ml/L |
| Vitamin K1 solution (see below) | 0.20 ml/L |
| Salt solution(DSMZ Salt solution): | |
| CaCl2 x 2 H2O | 0.25 g/L |
| MgSO4 x 7 H2O | 0.50 g/L |
| K2HPO4 | 1.00 g/L |
| KH2PO4 | 1.00 g/L |
| NaHCO3 | 10.00 g/L |
| NaCl | 2.00 g/L |
| Trace element solution(DSMZ Trace element solution): | |
| Nitrilotriacetic acid | 1.50 g/L |
| MgSO4 x 7 H2O | 3.00 g/L |
| MnSO4 x H2O | 0.50 g/L |
| NaCl | 1.00 g/L |
| FeSO4 x 7 H2O | 0.10 g/L |
| CoSO4 x 7 H2O | 0.18 g/L |
| CaCl2 x 2 H2O | 0.10 g/L |
| ZnSO4 x 7 H2O | 0.18 g/L |
| CuSO4 x 5 H2O | 0.01 g/L |
| KAl(SO4)2 x 12 H2O | 0.02 g/L |
| H3BO3 | 0.01 g/L |
| Na2MoO4 x 2 H2O | 0.01 g/L |
| NiCl2 x 6 H2O | 0.03 g/L |
| Na2SeO3 x 5 H2O | 0.30 mg/L |
| Vitamin solution: | |
| Biotin | 2.00 mg/L |
| Folic acid | 2.00 mg/L |
| Pyridoxine-HCl | 10.00 mg/L |
| Thiamine-HCl x 2 H2O | 5.00 mg/L |
| Riboflavin | 5.00 mg/L |
| Nicotinic acid | 5.00 mg/L |
| D-Ca-pantothenate | 5.00 mg/L |
| Vitamin B12 | 0.10 mg/L |
| p-Aminobenzoic acid | 5.00 mg/L |
| Lipoic acid | 5.00 mg/L |
| Haemin solution | |
| Dissolve 50 mg haemin in 1 ml 1 N NaOH; make up to 100 ml with distilled water. Store refrigerated. | |
| Vitamin K1 solution | |
| Dissolve 0.1 ml of vitamin K1 in 20 ml 95% ethanol and filter sterilize. Store refrigerated in a brown bottle. | |
| Others | |
| Na2SO4 | 0.1 g/L |
| FeSO4 | 0.05 g/L |
| (NH4)2SO4 | 0.09 g/L |

| Medium40. MPYG medium with SCFAs | |
|---|---|
| **Component** | **Concentration** |
| Trypticase peptone | 5.00 g/L |
| Peptone | 3.00 g/L |
| Peptone from soya | 2.00 g/L |
| Polypeptone | 1.00 g/L |
| Yeast extract | 10.00 g/L |
| Beef extract | 5.00 g/L |
| Glucose | 5.00 g/L |
| Tween 80 | 0.50 ml/L |
| Maltose | 0.50 g/L |
| Cellobiose | 0.50 g/L |
| Starch, soluble | 0.50 g/L |
| Glycerol | 0.50 ml /L |
| K2HPO4 | 2.00 g/L |
| Cysteine-HCl x H2O | 0.50 g/L |
| Na2S | 0.25 g/L |
| Resazurin | 1.00 mg/L |
| Salt solution (see below) | 40.00 ml/L |
| Trace element(see below) | 10.00 ml/L |
| Vitamin solution(see below) | 10.00 ml/L |
| Haemin solution (see below) | 10.00 ml/L |
| Vitamin K1 solution (see below) | 0.20 ml/L |
| Salt solution(DSMZ Salt solution): | |
| CaCl2 x 2 H2O | 0.25 g/L |
| MgSO4 x 7 H2O | 0.50 g/L |
| K2HPO4 | 1.00 g/L |
| KH2PO4 | 1.00 g/L |
| NaHCO3 | 10.00 g/L |
| NaCl | 2.00 g/L |
| Trace element solution(DSMZ Trace element solution): | |
| Nitrilotriacetic acid | 1.50 g/L |
| MgSO4 x 7 H2O | 3.00 g/L |
| MnSO4 x H2O | 0.50 g/L |
| NaCl | 1.00 g/L |
| FeSO4 x 7 H2O | 0.10 g/L |
| CoSO4 x 7 H2O | 0.18 g/L |
| CaCl2 x 2 H2O | 0.10 g/L |
| ZnSO4 x 7 H2O | 0.18 g/L |
| CuSO4 x 5 H2O | 0.01 g/L |
| KAl(SO4)2 x 12 H2O | 0.02 g/L |
| H3BO3 | 0.01 g/L |
| Na2MoO4 x 2 H2O | 0.01 g/L |
| NiCl2 x 6 H2O | 0.03 g/L |
| Na2SeO3 x 5 H2O | 0.30 mg/L |
| Vitamin solution: | |
| Biotin | 2.00 mg/L |
| Folic acid | 2.00 mg/L |
| Pyridoxine-HCl | 10.00 mg/L |
| Thiamine-HCl x 2 H2O | 5.00 mg/L |
| Riboflavin | 5.00 mg/L |
| Nicotinic acid | 5.00 mg/L |
| D-Ca-pantothenate | 5.00 mg/L |
| Vitamin B12 | 0.10 mg/L |
| p-Aminobenzoic acid | 5.00 mg/L |
| Lipoic acid | 5.00 mg/L |
| Haemin solution | |
| Dissolve 50 mg haemin in 1 ml 1 N NaOH; make up to 100 ml with distilled water. Store refrigerated. | |
| Vitamin K1 solution | |
| Dissolve 0.1 ml of vitamin K1 in 20 ml 95% ethanol and filter sterilize. Store refrigerated in a brown bottle. | |
| Others | |
| acetic acid | 33 mM |
| propionic acid | 8 mM |
| butyric acid | 4 mM |

| Medium41. GMM medium | | | |
|---|---|---|---|
| **Component** | **Amount/L** | **Concentration** | **Comments** |
| Tryptone Peptone | 2 g | 0.2% | |
| Yeast Extract | 1 g | 0.1% | |
| D-glucose | 0.4 g | 2.2 mM | |
| L-cysteine | 0.5 g | 3.2 mM | |
| Cellobiose | 1 g | 2.9 mM | |
| Maltose | 1 g | 2.8 mM | |
| Fructose | 1 g | 2.2 mM | |
| Meat Extract | 5 g | 0.5% | |
| KH2PO4 | 100 mL | 100 mM | 1M stock solution pH 7.2 |
| MgSO4-7$H_2$0 | 0.002 g | 0.008 mM | |
| NaHCO3 | 0.4 g | 4.8 mM | |
| NaCl2 | 0.08 g | 1.37 mM | |
| CaCl2 | 1 mL | 0.80% | 0.8g/100mL stock |
| Vitamin K (menadione) | 1 mL | 5.8 mM | 1 mg/mL stock solution |
| FeSO4 | 1 mL | 1.44 mM | 0.4 mg $FeSO_4$/mL stock solution |
| Histidine Hematin Solution | 1 mL | 0.1% | 1.2 mg hematin/mL in 0.2M histidine |
| Tween 80 | 2 mL | 0.05% | 25% stock solution |
| ATCC Vitamin Mix | 10 mL | 1% | |
| ATCC Trace Mineral Mix | 10 mL | 1% | |
| Acetic acid | 1.7 mL | 30 mM | |
| Isovaleric acid | 0.1 mL | 1 mM | |
| Propionic acid | 2 mL | 8 mM | |
| Butyric acid | 2 mL | 4 mM | |
| Resazurin | 4 mL | 4 mM | 0.25 mg/mL stock solution |
| Noble Agar | 12 g | 1.2% | |

| Medium42. GMM medium with mucin | | | |
|---|---|---|---|
| **Component** | **Amount/L** | **Concentration** | **Comments** |
| Tryptone Peptone | 2 g | 0.2% | |
| Yeast Extract | 1 g | 0.1% | |
| D-glucose | 0.4 g | 2.2 mM | |
| L-cysteine | 0.5 g | 3.2 mM | |
| Cellobiose | 1 g | 2.9 mM | |
| Maltose | 1 g | 2.8 mM | |
| Fructose | 1 g | 2.2 mM | |
| Meat Extract | 5 g | 0.5% | |
| KH2PO4 | 100 mL | 100 mM | 1M stock solution pH 7.2 |
| MgSO4-7$H_2$0 | 0.002 g | 0.008 mM | |
| NaHCO3 | 0.4 g | 4.8 mM | |
| NaCl2 | 0.08 g | 1.37 mM | |
| CaCl2 | 1 mL | 0.80% | 0.8g/100mL stock |
| Vitamin K (menadione) | 1 mL | 5.8 mM | 1 mg/mL stock solution |
| FeSO4 | 1 mL | 1.44 mM | 0.4 mg $FeSO_4$/mL stock solution |
| Histidine Hematin Solution | 1 mL | 0.1% | 1.2 mg hematin/mL in 0.2M histidine |
| Tween 80 | 2 mL | 0.05% | 25% stock solution |
| ATCC Vitamin Mix | 10 mL | 1% | |
| ATCC Trace Mineral Mix | 10 mL | 1% | |
| Acetic acid | 1.7 mL | 30 mM | |
| Isovaleric acid | 0.1 mL | 1 mM | |
| Propionic acid | 2 mL | 8 mM | |
| Butyric acid | 2 mL | 4 mM | |
| Resazurin | 4 mL | 4 mM | 0.25 mg/mL stock solution |
| Noble Agar | 12 g | 1.2% | |
| **Conponents that need to be added** | | | |
| **Component** | **Concentration** | | |
| Mucin | 4 g/L | | |

| Medium43. GMM medium with DSS | | | |
|---|---|---|---|
| **Component** | **Amount/L** | **Concentration** | **Comments** |
| Tryptone Peptone | 2 g | 0.2% | |
| Yeast Extract | 1 g | 0.1% | |
| D-glucose | 0.4 g | 2.2 mM | |
| L-cysteine | 0.5 g | 3.2 mM | |
| Cellobiose | 1 g | 2.9 mM | |
| Maltose | 1 g | 2.8 mM | |
| Fructose | 1 g | 2.2 mM | |
| Meat Extract | 5 g | 0.5% | |
| KH2PO4 | 100 mL | 100 mM | 1M stock solution pH 7.2 |
| MgSO4-7$H_20$ | 0.002 g | 0.008 mM | |
| NaHCO3 | 0.4 g | 4.8 mM | |
| NaCl2 | 0.08 g | 1.37 mM | |
| CaCl2 | 1 mL | 0.80% | 0.8g/100mL stock |
| Vitamin K (menadione) | 1 mL | 5.8 mM | 1 mg/mL stock solution |
| FeSO4 | 1 mL | 1.44 mM | 0.4 mg $FeSO_4$/mL stock solution |
| Histidine Hematin Solution | 1 mL | 0.1% | 1.2 mg hematin/mL in 0.2M histidine |
| Tween 80 | 2 mL | 0.05% | 25% stock solution |
| ATCC Vitamin Mix | 10 mL | 1% | |
| ATCC Trace Mineral Mix | 10 mL | 1% | |
| Acetic acid | 1.7 mL | 30 mM | |
| Isovaleric acid | 0.1 mL | 1 mM | |
| Propionic acid | 2 mL | 8 mM | |
| Butyric acid | 2 mL | 4 mM | |
| Resazurin | 4 mL | 4 mM | 0.25 mg/mL stock solution |
| Noble Agar | 12 g | 1.2% | |
| **Conponents that need to be added** | | | |
| **Component** | **Amount/L** | | |
| DSS | 1 g | | |

| Medium44. GMM medium with acetic acid | | | |
|---|---|---|---|
| **Component** | **Amount/L** | **Concentration** | **Comments** |
| Tryptone Peptone | 2 g | 0.2% | |
| Yeast Extract | 1 g | 0.1% | |
| D-glucose | 0.4 g | 2.2 mM | |
| L-cysteine | 0.5 g | 3.2 mM | |
| Cellobiose | 1 g | 2.9 mM | |
| Maltose | 1 g | 2.8 mM | |
| Fructose | 1 g | 2.2 mM | |
| Meat Extract | 5 g | 0.5% | |
| KH2PO4 | 100 mL | 100 mM | 1M stock solution pH 7.2 |
| MgSO4-7$H_20$ | 0.002 g | 0.008 mM | |
| NaHCO3 | 0.4 g | 4.8 mM | |
| NaCl2 | 0.08 g | 1.37 mM | |
| CaCl2 | 1 mL | 0.80% | 0.8g/100mL stock |
| Vitamin K (menadione) | 1 mL | 5.8 mM | 1 mg/mL stock solution |
| FeSO4 | 1 mL | 1.44 mM | 0.4 mg $FeSO_4$/mL stock solution |
| Histidine Hematin Solution | 1 mL | 0.1% | 1.2 mg hematin/mL in 0.2M histidine |
| Tween 80 | 2 mL | 0.05% | 25% stock solution |
| ATCC Vitamin Mix | 10 mL | 1% | |
| ATCC Trace Mineral Mix | 10 mL | 1% | |
| Acetic acid | 1.7 mL | 30 mM | |
| Isovaleric acid | 0.1 mL | 1 mM | |
| Propionic acid | 2 mL | 8 mM | |
| Butyric acid | 2 mL | 4 mM | |
| Resazurin | 4 mL | 4 mM | 0.25 mg/mL stock solution |
| Noble Agar | 12 g | 1.2% | |
| **Conponents that need to be added** | | | |
| **Component** | **Concentration** | | |
| acetic acid | 33 mM | | |

| Medium45. GMM medium with acetic acid, DSS and mucin | | | |
|---|---|---|---|
| **Component** | **Amount/L** | **Concentration** | **Comments** |
| Tryptone Peptone | 2 g | 0.2% | |
| Yeast Extract | 1 g | 0.1% | |
| D-glucose | 0.4 g | 2.2 mM | |
| L-cysteine | 0.5 g | 3.2 mM | |
| Cellobiose | 1 g | 2.9 mM | |
| Maltose | 1 g | 2.8 mM | |
| Fructose | 1 g | 2.2 mM | |
| Meat Extract | 5 g | 0.5% | |
| KH2PO4 | 100 mL | 100 mM | 1M stock solution pH 7.2 |
| MgSO4-7$H_2$0 | 0.002 g | 0.008 mM | |
| NaHCO3 | 0.4 g | 4.8 mM | |
| NaCl2 | 0.08 g | 1.37 mM | |
| CaCl2 | 1 mL | 0.80% | 0.8g/100mL stock |
| Vitamin K (menadione) | 1 mL | 5.8 mM | 1 mg/mL stock solution |
| FeSO4 | 1 mL | 1.44 mM | 0.4 mg $FeSO_4$/mL stock solution |
| Histidine Hematin Solution | 1 mL | 0.1% | 1.2 mg hematin/mL in 0.2M histidine |
| Tween 80 | 2 mL | 0.05% | 25% stock solution |
| ATCC Vitamin Mix | 10 mL | 1% | |
| ATCC Trace Mineral Mix | 10 mL | 1% | |
| Acetic acid | 1.7 mL | 30 mM | |
| Isovaleric acid | 0.1 mL | 1 mM | |
| Propionic acid | 2 mL | 8 mM | |
| Butyric acid | 2 mL | 4 mM | |
| Resazurin | 4 mL | 4 mM | 0.25 mg/mL stock solution |
| Noble Agar | 12 g | 1.2% | |
| **Conponents that need to be added** | | | |
| **Component** | **Concentration** | | |
| acetic acid | 33 mM | | |
| DSS | 1 g/L | | |
| Mucin | 4 g/L | | |

| Medium46. GMM medium with sulfate | | | |
|---|---|---|---|
| **Component** | **Amount/L** | **Concentration** | **Comments** |
| Tryptone Peptone | 2 g | 0.2% | |
| Yeast Extract | 1 g | 0.1% | |
| D-glucose | 0.4 g | 2.2 mM | |
| L-cysteine | 0.5 g | 3.2 mM | |
| Cellobiose | 1 g | 2.9 mM | |
| Maltose | 1 g | 2.8 mM | |
| Fructose | 1 g | 2.2 mM | |
| Meat Extract | 5 g | 0.5% | |
| KH2PO4 | 100 mL | 100 mM | 1M stock solution pH 7.2 |
| MgSO4-7$H_2$0 | 0.002 g | 0.008 mM | |
| NaHCO3 | 0.4 g | 4.8 mM | |
| NaCl2 | 0.08 g | 1.37 mM | |
| CaCl2 | 1 mL | 0.80% | 0.8g/100mL stock |
| Vitamin K (menadione) | 1 mL | 5.8 mM | 1 mg/mL stock solution |
| FeSO4 | 1 mL | 1.44 mM | 0.4 mg $FeSO_4$/mL stock solution |
| Histidine Hematin Solution | 1 mL | 0.1% | 1.2 mg hematin/mL in 0.2M histidine |
| Tween 80 | 2 mL | 0.05% | 25% stock solution |
| ATCC Vitamin Mix | 10 mL | 1% | |
| ATCC Trace Mineral Mix | 10 mL | 1% | |
| Acetic acid | 1.7 mL | 30 mM | |
| Isovaleric acid | 0.1 mL | 1 mM | |
| Propionic acid | 2 mL | 8 mM | |
| Butyric acid | 2 mL | 4 mM | |
| Resazurin | 4 mL | 4 mM | 0.25 mg/mL stock solution |
| Noble Agar | 12 g | 1.2% | |
| **Conponents that need to be added** | | | |
| **Component** | **Concentration** | | |
| Na2SO4 | 0.1 g/L | | |
| FeSO4 | 0.05 g/L | | |
| (NH4)2SO4 | 0.09 g/L | | |
